# Diesel exhaust particles disrupt mouse and human iPSC-derived microglial function and Amyloid-β clearance in Alzheimer’s disease models

**DOI:** 10.64898/2026.07.20.739493

**Authors:** Hong Yan, Yashvi Bhat, Polina Malahov, Angélica María Sabogal-Guáqueta, Teresa Mitchell-Garcia, Tingting Chen, Eugénie Genestant, Marko Iveša, Roberto Nebbia, Phoeja S. Gadjdjoe, Sohvi Ohtonen, Tarja Malm, Xavier Guillonneau, Martina Schmidt, Amalia M Dolga

## Abstract

Alzheimer’s disease (AD) is one of the most common neurodegenerative disorders, yet the environmental drivers that accelerate its progression remain poorly defined. Traffic-related air pollution is emerging as a modifiable AD risk factor, but how inhaled particles perturb microglial clearance of amyloid beta (Aβ) is unknown. Microglia are the principal Aβ-clearing phagocytes of the brain. Here, we showed that exposure of primary mouse microglia and human induced pluripotent stem cell-derived microglia (iMGLs) to 3–100 µg/mL diesel exhaust particles (DEP) disrupted microglial homeostasis, induced morphological abnormalities, increased reactive oxygen species, impaired lysosomal degradation, and led to a concentration-dependent loss of phagocytic capacity. Importantly, DEP markedly reduces Aβ uptake in both species. Transcriptomic profiling revealed a DEP-induced, non-canonical state characterized by metabolic reprogramming, broad suppression of inflammatory pathways, antigen-presentation, chemokine, and species-specific remodeling during subsequent Aβ challenge, including defective chemotaxis, cell cycle, and cytoskeletal signatures. These data show that DEP profoundly alters microglial transcriptional and metabolic states, leading to impaired Aβ clearance, which could, thereby, further contribute to AD progression.

## 1. Introduction

Alzheimer’s disease (AD) is one of the most prevalent degenerative neurological disorders, and is highly associated with aging [1]. In 2019, 57.4 million people were afflicted with dementia; and this number is predicted to increase to 152.8 million in 2050 [2]. The main features of primary AD pathology are represented by the presence of extracellular amyloid-beta (Aβ) deposits and intracellular neurofibrillary tangles, as a result of tau hyperphosphorylation [3]. The process of Aβ deposition is considered to be toxic to neurons, as during this process there is an increase in oxidative stress and proinflammatory responses [1, 4]. The equilibrium between the continuous generation of Aβ and its effective clearance is crucial for maintaining Aβ homeostasis and extensive research has focused on strategies on targeting the toxic aggregation and misfolded assemblies [5]. Although the most studied risk factors for AD were linked to the familial forms of AD [6], substantial evidence indicates that external environmental factors could significantly influence the development and progression of AD [7–9].

Air pollution can be defined as a complex mixture of particulate matter (PM), (micro)plastics, organic compounds, gases, and metals. PM stands out as a major health hazard due to its link with an increased onset of pulmonary, cardiovascular, central nervous system (CNS), or immune diseases [10, 11]. Recent research has shown that fine (PM_2.5_; diameter ≤ 2.5 µm) and ultrafine particles (PM_0.1_; diameter ≤ 0.1 µm) are considered the most important risk factors affecting health due to their ability to penetrate deep into the tissue [12–14]. PM can deposit in the alveolar lung regions, infiltrate the lung epithelium, and ultimately reaching the systemic circulation. Following systemic absorption, the circulating particles subsequently reach the blood-brain barrier (BBB), where they may potentially infiltrate the BBB and translocate into the brain tissue [15, 16].

A number of clinical, epidemiological, and experimental studies have documented the impact of air pollution on the development of AD, with a particular focus on its effects on the CNS [17–19]. Recently it was reported that urban dogs chronically exposed to PM in Mexico City exhibited impairment of the BBB and extracellular deposition of Aβ peptide fibrils in subcortical and cortical structures, which indicated that air pollution may play a critical role in initiating AD early in life Calderón-Garcidueñas, et al. [20]. A study in young people (≤30 years) in Mexico City showed that PM_2.5_ could lead to early neuroinflammation and neurodegeneration in children [21, 22]. Exposure to PM_2.5_ prenatally resulted in long-lasting memory deficits in male mice offspring, potentially mediated by DNA methylation-associated inflammatory responses [23]. A two-week exposure to PM_0.1_ in aged 3xTgAD mice at concentrations relevant to human exposure resulted in memory deficits [24]. Moreover, exposure to PM_2.5_ resulted in neuroinflammation and neurodegeneration, driven by the activation of astrocytes and microglia in 3D brain models [25].

Microglia are resident immune cells in the CNS, accounting for 5-15% of the adult brain cells [26]. They originate in the yolk sac as primitive hematopoietic stem cells during early development, then migrate to the CNS, where they increase in number and mature into microglia [27]. Microglia can quickly sense diverse disturbances in the CNS, ranging from internal signals such as cellular debris and misfolded proteins (e.g., Aβ deposits) to external insults including pathogens and air pollutants [26, 28]. PM exposure may influence the function of microglia, which in turn can affect an individual’s susceptibility to AD, as microglial dysfunction poses a significant risk for the progression of AD pathology [29, 30]. In response to external stimuli or stress, microglial cells can display a spectrum of different morphological and functional states [31, 32]. Homeostatic microglia have a more branched morphology, elongated processes, and smaller cell bodies during the surveillance of their microenvironment, whereas responsive microglia change to a more circular, amoeboid morphology, retracted processes, and enlarged cell bodies [33]. Diesel exhaust particles (DEP) application through intranasal administration resulted in an increase microglia response resulting in the mouse hippocampus [34]. Long-term changes in microglial morphology and neuron-glia interactions were also demonstrated in mouse brain following prenatal DEP oropharyngeal aspiration exposure, a model usually used to assess lung inflammation and toxicity [35].

Phagocytosis is a key microglial function and, along with clathrin-mediated endocytosis and macropinocytosis, represents distinct form of endocytosis [36]. Microglia engulf foreign particles >0.5 µm through phagocytosis [37]. Internalized cargo affects early and late endosomes [38], leading to progressive acidification and degradation [39]. Markers such as early endosome antigen 1 (EEA1) and lysosomal-associated membrane protein 1 (LAMP1) are commonly used to assess endosomal and lysosomal function [40, 41]. Efficient lysosomal acidification is essential for phagocytosis and autophagy-mediated degradation [42, 43]. As the final stage of clearance, lysosomal pH and enzymatic activity critically regulate cargo degradation, thereby influencing microglial activation in health and disease [44, 45]. Defective lysosomal acidification disrupts microglial homeostasis, promoting neuroinflammation and neurodegeneration [46]. In AD, microglia exhibit dual roles in Aβ clearance: early responses promote uptake, while chronic stimulation impairs Aβ degradation and exacerbates pathology [47–50].

Whilst most insights into microglial biology stem from rodent models, accumulating evidence highlights significant species-specific differences between mouse and human microglia [51]. Human microglia display distinct transcriptional signatures, developmental trajectories, and inflammatory responses compared to their murine counterparts, which may influence phagocytosis, lysosomal function, and cytokine release. These differences are particularly relevant for neurodegenerative diseases such as AD, as human microglia could exhibit altered interactions with Aβ and provide an inflammatory profile to the surrounding cells. Therefore, including human microglia as a model system is critical to better understand disease mechanisms and to ensure translational value when interpreting microglial responses across species.

In this study, we assessed whether DEP can contribute to the AD pathology and which pathways are involved in both mouse microglia and human iPSC-derived microglia. We investigated the transcriptome profile together with the potential phenotypic alterations, phagocytosis, lysosomal function, reactive oxygen species (ROS) production in microglia exposed to DEP. We studied these processes in human iPSC-derived microglia to demonstrate the translational potential of DEP exposure, and to uncover cell susceptibility to AD.

## 2. Methods

### 2.1 Animals

C57BL/6 J mice were obtained from the central animal laboratory at University of Groningen. All animal care and use complied with Dutch standards and guidelines (Protocol 2215899-01-002). All experimental procedures and protocols were approved by the University of Groningen Committee for Animal Experimentation.

### 2.2 Isolation of primary mouse microglia

Primary mouse microglia were isolated from day 0 to day 3 of C57BL/6 pups as previously described [51, 52]. Cortices from pups were dissected, stripped of meninges, and dissociated with 0.25% Trypsin and DNAse for 25 min. The cell suspension was seeded onto T75 tissue culture flasks and cultured in microglia medium (MM)-Dulbecco’s Modified Eagle Medium (DMEM; 42430025; Gibco, Thermo Fisher Scientific) supplemented with 10% (v/v) fetal bovine serum (FBS; SV3016.03; Cytiva HyClone, Fisher Scientific), 2% (v/v) Pen-Strep (15070063; 5 × 10^3^ U/mL; Gibco, Thermo Fisher Scientific), and 1% (v/v) sodium pyruvate (11360070; 100 mM; Gibco, Thermo Fisher Scientific) - at 37 °C and 5% CO_2_ for 10 - 14 days to grow a confluent layer of astrocyte. After 4 - 6 days, flasks were shaken at 37 °C and 150 rpm for 60 min, followed by collecting microglia from the supernatant by centrifuging at 230 rcf for 12 min at room temperature (RT). After that, microglia were maintained in a mixture of 50% astrocyte-conditioned medium and 50% fresh MM. For all experiments, only primary microglial cells from the first, second, and third collections were used.

### 2.3 iPSC microglia generation

The induced pluripotent stem cell (iPSC) line purchased from Gibco (A18945) was used for the microglia differentiation. The iPSC colonies were cultured and expanded onto Matrigel (354277, Corning) in 6-well plates in E8 flex medium (A2858501, Gibco) supplemented with 100 U/mL Pen-Strep and E8 flex supplement. Once the cells were confluent, iPSC colonies were passed every 3-4 days using enzymatic detachment with RelesR (100-0483, STEMCELL Tech) for three minutes at 37 °C, 5% CO_2_, and re-plated in E8 flex medium.

Microglia differentiation from iPSCs was based on modified protocols [51]. Cells at 70 to 80% confluent in 6-well plates were used for day 0 of differentiation. Cells were detached with Accutase (A6964, Sigma Aldrich) and incubated at 37 °C for 5 min. Detached cells were resuspended with 1× PBS, then transferred to a Falcon tube and centrifuged for 5 min at 300 rcf. After aspirating the supernatant, the cell pellet was resuspended in differentiation medium 1 comprising of E8 flex supplemented with 100 U/mL Pen-Strep, E8 flex supplement, 50 ng/mL BMP-4 (120-05, Peprotech), 50 ng/mL VEGF (100-20, Peprotech), 20 ng/mL SCF (300-07, Peprotech), and 10 µM Rock Inhibitor (688001-500UG, MERCK). Cells were seeded at a density of 10,000–15,000 cells/well in a Nuncleon 96-well U-bottom sphera-coated plate (174929, Thermofisher). The plate was centrifuged at 100 rcf for 5 min to ensure clustering of the cells at the bottom of the well, and then incubated at 37 °C, 5% CO_2_. From day 1 to day 3, 75% of medium was replaced with 100 µL/well fresh differentiation medium 1. On day 4, all the embryoid bodies (EBs) were harvested, resuspended in 12 mL differentiation medium 2 consisting of XVIVO 15 medium (BE02-060Q, LONZA), 2 mM Glutamax (35050061, Gibco), 100 U/mL Pen-Strep and 0.055 mM 2-mercaptoethanol (31350010, Thermofisher) with 50 ng/mL SCF, 50 ng/mL M-CSF (300-25, Peprotech), 50 ng/mL IL3 (200-03, Peprotech), 50 ng/mL FLT3 (300-19, Peprotech) and 5 ng/mL TPO (300-18, Peprotech). The EBs in this medium were transferred to a T75 flask. On day 8, the old medium was changed to 12 mL of fresh differentiation medium 2. The old medium was refreshed also on days 11 and 18 with 12 mL fresh differentiation medium 3 consisting of XVIVO 15, 2 mM Glutamax, 100 U/mL Pen-Strep, 0.055 mM 2-mercaptoethanol, supplemented with 50 ng/mL FLT-3, 50 ng/mL M-CSF, and 25 ng/mL of GM-CSF (300-03, Peprotech). On day 25, the medium in the flask was collected, then passed through a 40 µm cell strainer and centrifuged at 300 rcf for 5 min. The cells in the pellet at this stage are called microglia progenitors. These were then resuspended in maturation medium comprising Gibco Advanced DMEM/F12 supplemented with 5 µg/mL N-acetylcysteine (A0737, Sigma-Aldrich), 400 µM 1-Thioglycerol (M6145, Sigma-Aldrich), 1 µg/mL heparan sulfate (H7640, Sigma-Aldrich), 1% Glutamax, 1% NEAA (11140-050, Gibco), 1% Pen-Strep, 2% B27 without vitamin A (12587010, Gibco), and 0.5% N2 (17502-048, Gibco). Growth factors: 100 ng/mL interleukin-34 (200-34, Peprotech), 25 ng/mL M-CSF, 25 ng/mL CX3CL1 (300-31, Peprotech), 25 ng/mL TGF-β-1 (100-21C, Peprotech), and eventually formed iPSC-derived microglia-like (iMGL) cells.

### 2.4 Preparation of diesel exhaust particles

The standard reference material (SRM) 2975 diesel exhaust particle (DEP) was obtained from the National Institute of Standards and Technology (1333-86-4, Industrial Forklift) and prepared as previously described [53, 54]. Briefly, dispersed suspensions of DEP were prepared by sonication in PBS for 30 min in a cooling water bath (06524, Branson 2510; Gemini b.v.) followed by probe sonication (20 kHz, 130 W; Ultrasonic Processor, Sonics Vibra Cell) for 30 sec. Then 0.1% (v/v) Tween 80 (9005-65-6, Sigma-Aldrich) was added to ensure proper resuspension of particles [55]. DEP was used at 3, 10, 30, and 100 µg/mL concentrations. Controls were prepared in PBS with 0.1% (v/v) Tween 80, following the same steps as DEP.

### 2.5 Microglial morphology analysis

Primary mouse microglia were seeded onto 96-well flat-bottom plates at a density of 15,000 cells/well 48 h before treatment. Cells were then exposed to the indicated concentrations of DEP. IncuCyte S3 Live-Cell Analysis Instrument (4647, Essen Bioscience) was used to monitor morphological changes at 10× and 20× magnification for 24 h. The phase eccentricity object average was obtained using the IncuCyte S3. Eccentricity defines the range of circularity of an object, ranging from 0 to 1. For a perfect circle, the eccentricity is exactly 0. In addition, for roundness analysis, the processed images were automatically thresholded using the maximum entropy algorithm, and binary masks were generated. For each image, roundness was measured using the Fiji/ImageJ built-in measurement function.

In addition, to analyze single isolated microglia, higher-resolution images were captured by Nikon Inverted Research Fluorescence Microscope Eclipse Ti2-E. Primary microglia were seeded onto 8-well µ-slide plates (Ibidi GmbH) at a density of 30,000 cells/well 48 h before treatment. After exposure to DEP at concentrations of 3, 10, and 30 µg/mL, single-cell morphology was analyzed using fractal analysis with the FracLac plugin of ImageJ/Fiji software (https://imagej.net/ij/plugins/fraclac/fraclac.html) [56]. Briefly, FFT bandpass filter, unsharp mask, and close were consistently used before converting all photomicrographs to the binary format. Four microglia were randomly chosen for fractal analysis (using predefined positions) within the photomicrograph (4 photomicrographs per condition) for a total of 16 cells analyzed per condition. The extra formations adjacent to and enclosing each cell were removed from the analysis by manually deleting them using the ImageJ/Fiji software. After that, the binary images were transformed into outlines. Span ratio and circularity were quantified using the FracLac-generated “convex hull” which circumscribes each microglia outline with a polygon and a circle that bounds the convex hull. Span ratio can be defined as the ratio of the convex hull ellipse’s longest length to the longest width to describe the elongation of the cell outline. Circularity is used to describe the shape of the cell outline and is the comparison of the convex hull area to the convex hull perimeter, ranging from 0 to 1. For a perfect circle, the circularity is exactly 1.

### 2.6 Phagocytosis assay

The phagocytic capacity of mouse and iMGL cells was measured using pHrodo red *E. coli* bioparticles (P35361, Thermo Fischer) with IncuCyte S3 Live-Cell Analysis Instrument (4647, Essen Bioscience) [51, 57]. Mouse and iMGL cells were seeded onto 96-well flat-bottom plates at a density of 15,000 cells/well 48 h before treatment. For pre-exposure, mouse microglia were incubated with DEP at concentrations of 3, 10, and 30 µg/mL for 24 h, and iMGL cells were exposed to 20 µg/mL of DEP for 24 h. Then, cells were incubated with 1 µg/0.1 mL pHrodo red bioparticles and monitored in IncuCyte for a minimum of 24 h. For co-exposure experiments, mouse microglia were treated with the combination of 1 µg/0.1 mL pHrodo red bioparticles and DEP at 3, 10, and 30 µg/mL, respectively. Cells were imaged with phase contrast and red fluorescent channels at 20× magnification every hour for 24 h. Utilizing the Incucyte Software, the integrated fluorescence intensity was measured following the exclusion of background fluorescence via adaptive segmentation thresholding.

### 2.7 Lysosomal function

#### Incucyte imaging

Lysosomal activity was measured as described before [58]. Primary mouse microglia were seeded onto 96-well flat-bottom plates at a density of 15,000 cells/well 48 h before treatment. Ammonium chloride (NH_4_Cl) was used as a negative control as it causes a transient decrease in the lysosomal pH. DMEM medium (not supplemented with FBS) was used as positive control because in primary mouse microglia it induced autophagy and increased lysosomal staining. Cells were stained with Lysoview 540 reagent (70061, Biotium, 1×) for 30 min, followed by taking the first image as a timepoint zero (T0) to evaluate the basal lysosomal activity per well. Then, the cells were exposed to DEP at concentrations of 3, 10, and 30 µg/mL. Phase-contrast and red fluorescent images were imaged at 10× magnification every hour for 24 h. The integrated fluorescence intensity of the red channel was measured following the exclusion of background fluorescence via adaptive segmentation thresholding and normalized to the T0 measurement.

#### Live cell imaging

Primary mouse microglia were seeded onto 8-well µ-slide plates (Ibidi GmbH) at a density of 30,000 cells/well 48 h before treatment. Cells were stained with a combination of Lysoview 488 (70067, Biotium, 1×) and Lysoview 540 (70061, Biotium, 1×) for 30 min. Then, the cells were exposed to DEP at concentrations of 3, 10, and 30 µg/mL. Following an incubation time of 2 h, fluorescence was detected at Ex/Em = 496/526 (Lysoview 488) and 540/634 (Lysoview 540) using a Nikon Inverted Research Fluorescence Microscope ECLIPSE Ti2-E at a constant exposure of 100 ms. Images of each well were captured at a magnification of 60×.

### 2.8 Immunofluorescence

Primary mouse microglia were seeded onto removable 12-well chamber slides (Ibidi GmbH) at a density of 20,000 cells/well 48 h before treatment. Then, cells were exposed to DEP at concentrations of 3, 10, and 30 µg/mL for 24 h. For LAMP1 or EEA1 staining, cells were fixed with 4% paraformaldehyde solution (pH 7.4) at room temperature (RT) for 15 min, followed by incubation with blocking and permeabilization buffer (3% BSA, 0.1% TritonX-100 in PBS) at RT for 30 min. The cells were then incubated with LAMP1 (1:400; AB2971-25UG, Sigma-Aldrich) and EEA1 antibody (1:200; 610457, BD Transduction Laboratories™) in the dark at 4 °C overnight. After a thorough wash in PBS buffer, the cells were incubated with Alexa Fluor 488-conjugated donkey anti-rabbit IgG (H+L) secondary antibody (A-21206; Invitrogen, Thermo Fisher Scientific, the Netherlands) and Alexa Fluor 568-conjugated donkey anti-mouse IgG (H+L) secondary antibody (A-10037; Invitrogen, Thermo Fisher Scientific, the Netherlands) for 2 h. Incubations with only secondary antibodies were performed simultaneously as controls for the immunostaining procedure. After removing the silicone gasket, coverslips were mounted onto slides by antifade mounting medium Fluoroshield™ with DAPI (F6057-20ML, Sigma-Aldrich). Fluorescence was detected at Ex/Em = 490/525 nm for LAMP1 and Ex/Em = 579/603 nm for EEA1 on a Nikon Inverted Research Fluorescence Microscope ECLIPSE Ti2-E. At least 15 images per well were taken at 60× magnification.

### 2.9 ROS assay

The intracellular accumulation of reactive oxygen species (ROS) was assessed using CellROX Deep Red reagent (C10422, Invitrogen) and IncuCyte live-cell imaging [57, 59]. CellROX Deep Red is a fluorescent ROS sensor that does not fluoresce in the reduced state but produces bright near-infrared fluorescence when oxidized by ROS. Primary mouse microglia were seeded onto 96-well flat-bottom plates at a density of 15,000 cells/well 48 h before treatment. Cells were incubated with 1.25 µM CellROX Deep Red for 30 min, and then exposed to DEP at concentrations of 3, 10, and 30 µg/mL. Phase contrast and red fluorescence images were acquired every hour for 24 h at 10× magnification. The integrated fluorescence intensity of the red channel was measured following the exclusion of background fluorescence via adaptive segmentation thresholding.

### 2.10 Clearance of Amyloid beta (Aβ)

Human Aβ (1–42) labeled with HiLyte™ Fluor 488 (AS-60479-01, ex/em 503/528 nm) was purchased from AnaSpec Inc and prepared according to the manufacturer’s instructions. This was a mixture of monomers and amorphous aggregates of various sizes and shapes. Mouse and iMGL cells were seeded onto 8-well ibidi plates at a density of 20,000 cells per well 48 h before treatment. Then, cells were exposed to DEP at concentrations of 3, 10, and 30 µg/mL at 37 °C and 5% CO_2_ for 24 h. After pre-treatment, the cells were incubated using 488-labeled Aβ, and images were acquired using IncuCyte live-cell imaging continually every hour for 24 h, and a Nikon Eclipse Ti camera after 4 h.

### 2.11 RNA extraction for bulk sequencing

For RNA collection, mouse and iMGL cells were seeded in 6-well plates at a density of 450,000 cells/well, 48 h before treatment. Cells were exposed to 10 µg/mL diesel exhaust particles (DEP) for 4 or 24 h. In the 24 h DEP condition, microglia were subsequently treated with amyloid-β (Aβ) for 4 h. After treatment, cells were washed twice with ice-cold PBS, and the dried plates were immediately snap-frozen on dry ice.

RNA extractions were performed using NucleoSpin® RNA isolation kits (740990 and 740962, Macherey-Nagel). RNA sequencing libraries were constructed from 250 ng total RNA using a modified TruSeq RNA Sample preparation kit protocol. Pass-filtered reads (using Trimmomatic) were mapped using HiSAT2 and aligned to human reference genome GRCh38 and mouse reference genome GRCm39 [60].

The count table of the gene features was obtained using HTSeq. Normalization and differential expression analysis values were computed using DESeq2 [61]. TPMs were determined using Libinorm using the htseq mode [62].

### 2.12 RNAseq analysis

Fastq files obtained from the sequencing facility were aligned using STAR (v2.7.9a) against the organism reference genome, with option “--quantMode GeneCounts” to extract the raw counts for each gene, then all count files were concatenated into a single file. A sample file was created with the sample data including the conditions (DEP4, DEP24, DEP Aβ4) and the replicate numbers. The count file and the sample file were loaded in an in-house R Shiny application EYE DV seq. We added 1 to all counts in the count file to avoid any 0 read count errors. We then removed the genes having a total count < 10 across all samples with an FDR rate of 10%. Finally, the DESeq2 (v1.40.2) analysis was performed. The results for read count were then filtered for significant genes (Padj < 0.05). PCA analysis was performed using pcaExplorer (v2.9.6). Further filtering was applied for an enrichment analysis (Padj < 0.05,-3 < log2FC < 3). The enrichment analysis was performed to study the implicated pathways, using the GO database (using the Bioconductor packages and ‘clusterProfiler’ (v 4.8.1)).

### 2.13 Statistical analysis

All experiments were conducted in triplicate wells and replicated three times. One-way ANOVA with Tukey’s post-hoc test was used. All groups were compared to the control (solvent medium) condition. Statistical analyses and graphing were performed using GraphPad Prism 9 (GraphPad Software), and *p*-values below 0.05 were considered significant.

## 3. Results

### 3.1. DEP exposure alters primary mouse microglia shape and length

Microglia maintain brain homeostasis by continuously sensing and responding to changes in their microenvironment [63]. Upon stimulation, microglial morphology and motility undergo substantial alterations [32]. In this study, we used primary mouse microglia and human iMGLs as they reflect well the physiological state of brain microglia compared to immortalized microglial cell lines [64, 65] [66]. Live-cell imaging experiments were performed to assess primary mouse microglial responses following 24-hour exposure to various concentrations of DEP (3–100 µg/mL). We used these concentrations based on previous studies [54, 67]. As shown in **Figure 1A** and **supplementary S1**, 30 and 100 µg/mL of DEP exposure resulted in visibly large cell bodies, and the analysis of the cell morphology indicated a significant increase towards cell roundness form, with an amoeboid morphology. As our aim was to use DEP concentrations that are not oversaturating the microglial phagocytic capacity, subsequent experiments were conducted using lower DEP concentrations (3–30 µg/mL).

**Figure 1.**
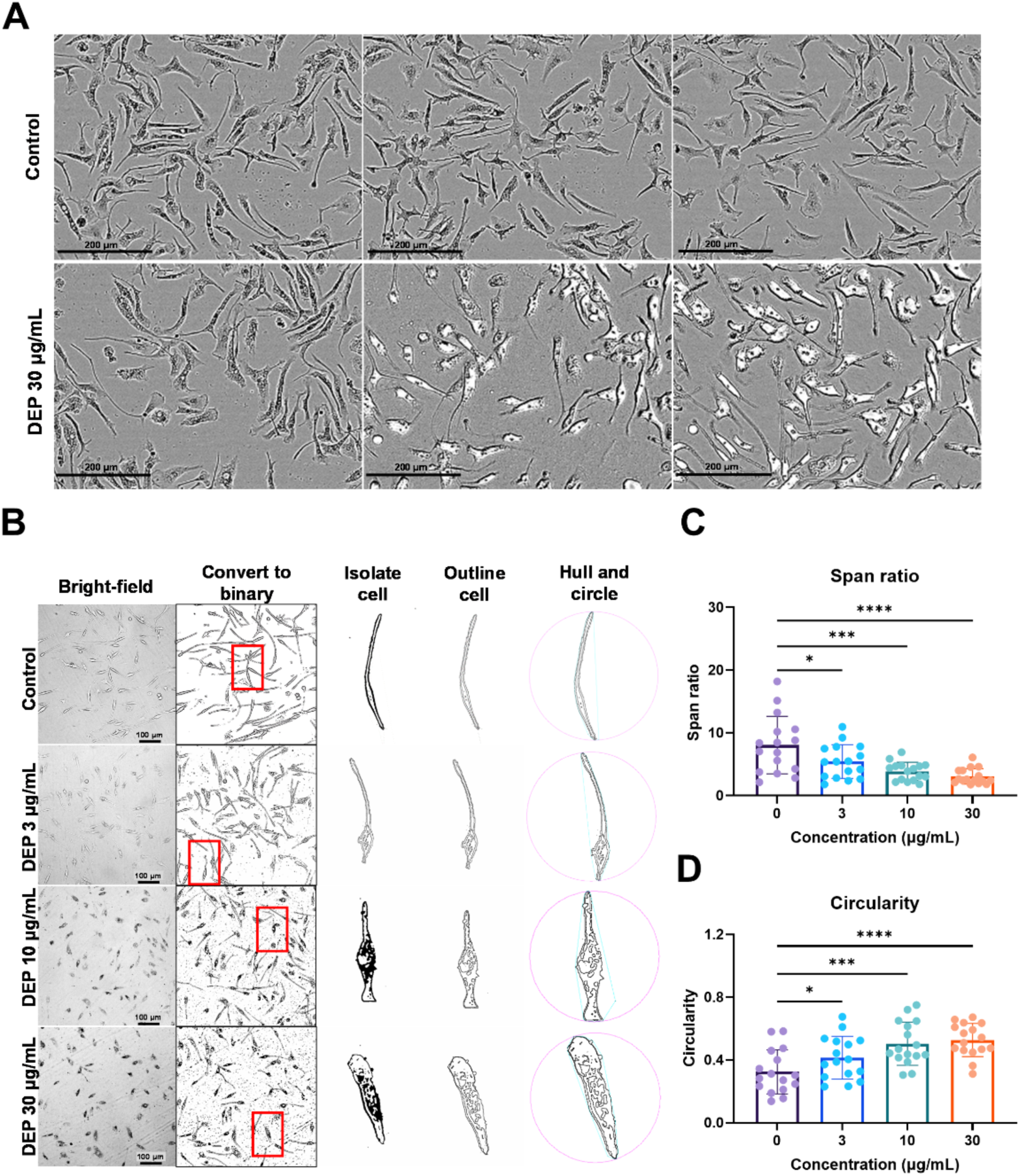
Morphological changes in mouse microglia following DEP exposure. (**A**) Representative phase-contrast images of primary microglial cells treated with solvent control or 30 µg/mL DEP at 0, 12, and 24 h. Scale bar = 200 µm. (**B**) Brightfield photomicrographs and cell outlines of unstained microglial cells. Scale bar = 100 µm. (**C, D**) Quantification of span ratio (**C**) and circularity (**D**) of microglia exposure to 0, 10, and 30 µg/mL of DEP. All data are mean ± SD with one-way ANOVA. \**p* < 0.05, \*\*\**p* < 0.001, \*\*\*\**p* < 0.0001. 16 photomicrographs per condition were analysed per experiment. Each experiment was repeated three times.

Recent studies have shown that single-cell analytical approaches offer greater precision in assessing microglial morphology than analyses based on whole-field photomicrographs of cell population [68]. Accordingly, we applied a single-cell analysis workflow using photomicrographs to quantify span ratio and circularity. Single microglial cells were digitally isolated, converted into a binary format, outlined, and the morphometric measurements were performed using the FracLac plugin for ImageJ (**Figure 1B**). We analyzed circularity and span ratio, as key descriptors of microglial roundness and elongation. These two measurements are particularly relevant to microglial roundness as they are directly linked to the cell shape/length. As shown in **Figures 1C and D**, DEP exposure induced a statistically significant increase in circularity and a decrease in span ratio in a concentration-dependent way. These data demonstrate that DEP exposure drives a shift toward an activated morphological state, typically reflected by the cell process retraction and increased cell body roundness.

### 3.2. DEP impairs phagocytosis in microglial cells

Phagocytosis of cell debris, neuronal synapses, and apoptotic cells represent key functions of microglia [45, 69]. To determine whether DEP exposure impacts the microglial ability of phagocytosis, we assess the real-time uptake of pHrodo red *E.coli* bioparticles using live imaging. Primary mouse microglial cells were pre-incubated with DEP at concentrations of 3, 10, and 30 µg/mL for 24 hours, followed by incubation with pHrodo red *E.coli* bioparticles. As shown in **Figure 2A and Supplementary figure S2**, red-channel fluorescence images revealed a progressive decrease in microglial fluorescence intensity with increasing concentrations of DEP. Time-course analysis of integrated red fluorescence (**Figure 2B**) confirmed a concentration-dependent reduction in bioparticle uptake, indicating a gradual decline in the phagocytic capacity of microglial cells over time. Quantification at 24-hour (**Figure 2C**) showed that all tested DEP concentrations significantly reduced the uptake of pHrodo-labeled particles compared with solvent-treated controls.

**Figure 2.**
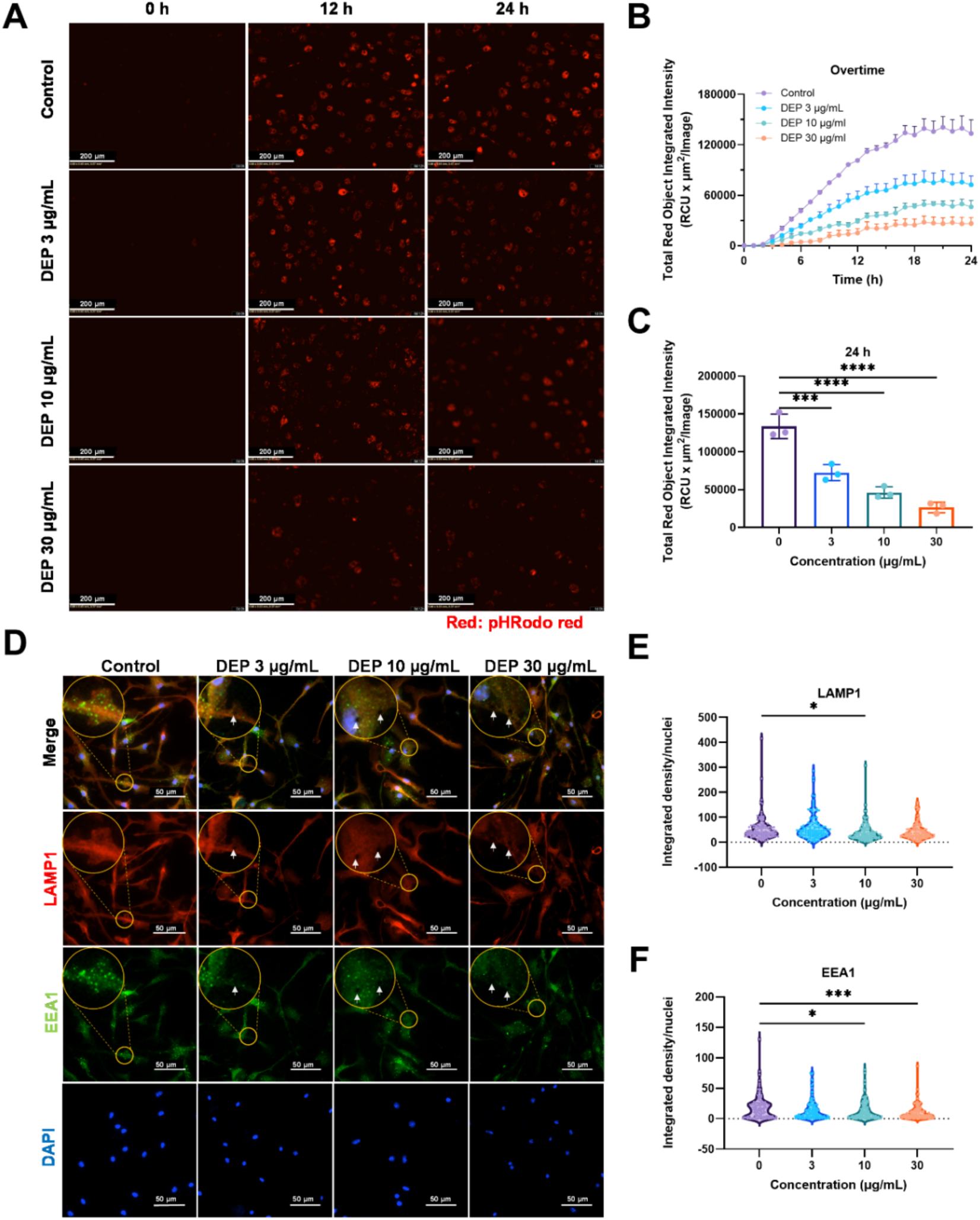
DEP impairs phagocytosis in mouse microglia. (**A**) Representative red fluorescence images of primary microglial cells showing phagocytosis of pHrodo red *E.coli* bioparticles after pretreatment with solvent control, 3, 10, or 30 µg/mL DEP at 0, 12, and 24 h. Scale bar = 200 µm. (**B**) Time-curve of integrated red fluorescence intensity in microglia pretreated with 3, 10, and 30 µg/mL DEP. (**C**) Bar graph for the integrated fluorescence intensity of pHrodo red *E.coli* bioparticles at 24 h after indicated exposure. (**D**) Representative images of primary mouse microglia treated with 3, 10, and 30 µg/mL DEP for 24 h then stained with EEA1 (green), LAMP1 (red), and DAPI (nuclei, blue). White arrows indicated the area with DEP. Scale bar = 50 µm. **(E, F)** Bar graphs showing integrated fluorescence intensity normalized on the number of nuclei of LAMP1 (**E**) and EEA (**F**) in microglia after the indicated treatment. Individual data points represent results from four independent experiments. All data are mean ± SD with one-way ANOVA. \**p* < 0.05, \*\*\**p* < 0.001, \*\*\*\**p* < 0.0001. n=3 independent experiments with 3 wells per condition.

To investigate whether DEP-induced impairment of phagocytosis occurs at specific stages of phagosome maturation, we performed double immunofluorescence staining for EEA1 (early endosomes) and LAMP1 (late endosomes/lysosomes). Microglial cells were exposed to increasing concentrations of DEP (3, 10, and 30 µg/mL) and analyzed for changes in these endosomal markers. As shown in **Figure 2D**, DEP exposure resulted in a reduction of both LAMP1 and EEA1 fluorescence intensities, particularly in the regions containing engulfed DEP (indicated by white arrows). At 10 µg/mL DEP, the fluorescence intensities of both markers decreased significantly (**Figures 2E and 2F**). These results indicate that DEP affects endosomal and lysosomal compartments, suggesting that the impaired phagocytic capacity of DEP-treated microglia could be associated with disturbances in endosomal trafficking and phagosome maturation.

### 3.3. DEP induces lysosomal dysfunction in primary mouse microglia

Our initial results showed a clear concentration-dependent response of microglia to DEP exposure, accompanied by impaired phagocytic activity. Given the critical importance of lysosomal function in regulating both autophagy and phagocytosis in microglia [43], we next assessed whether DEP disrupts lysosomal homeostasis. The analysis of the DEP exposed microglia showed that increasing DEP concentrations reduced Lysoview 540 (a pH-sensitive probe) fluorescence in primary mouse microglial cells, indicating progressive lysosomal alkalinization (**Figure 3A and B**), with decreased fluorescence intensities even after short time exposure (2 hours) to DEP (**Figure 3C**). These results demonstrated that DEP induces a progressive loss of lysosomal acidity.

**Figure 3.**
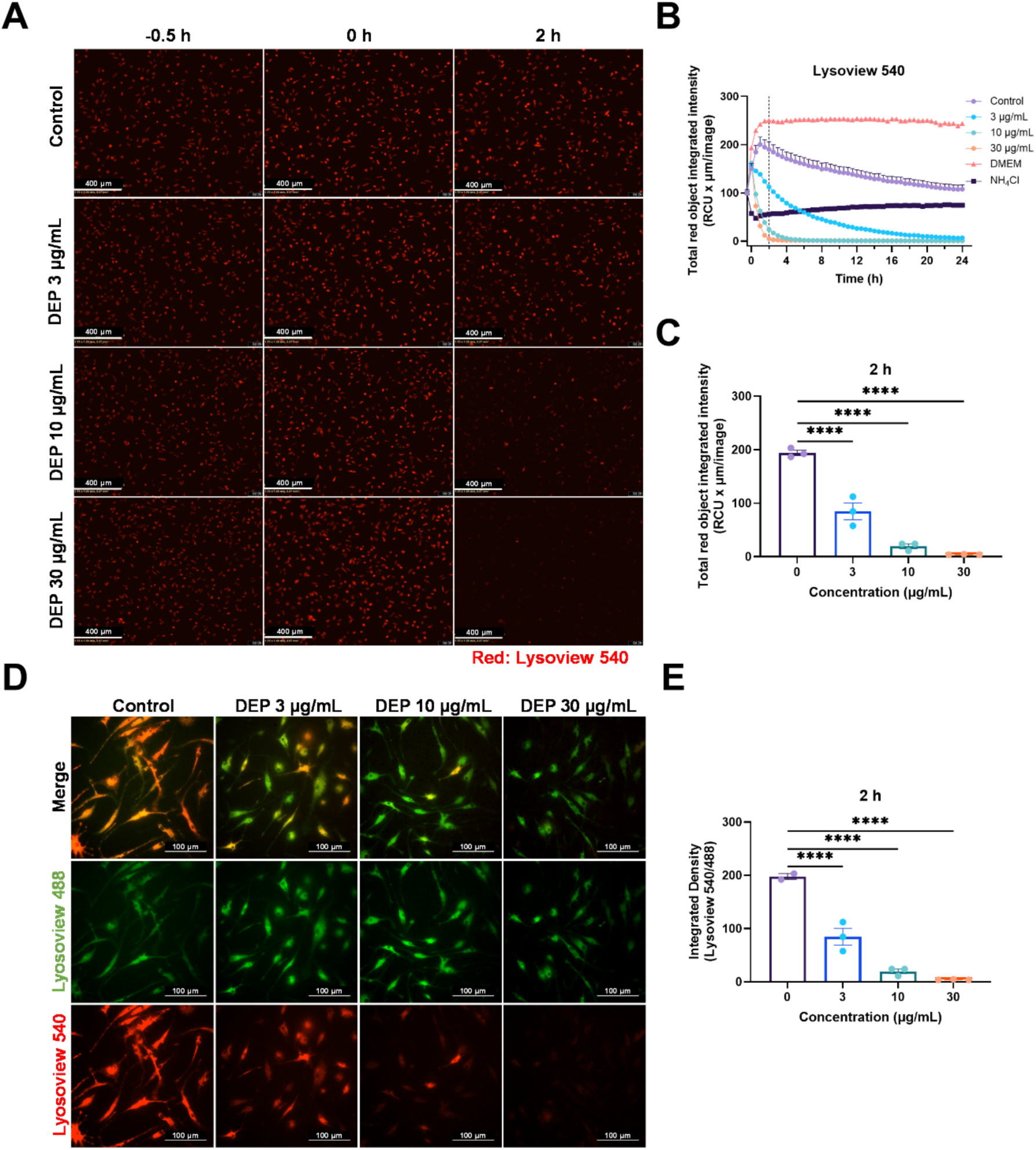
DEP causes lysosomal alkalization in primary mouse microglia cells. (**A**) Representative images of primary microglial cells with phase contrast and red fluorescent channel showing stained lysosomes of microglial cells treated with 3, 10, and 30 µg/mL DEP at-0.5 (t0), 0, and 2 h. Scale bar = 400 µm. (**B**) Time-curve of integrated red fluorescence intensity for 3, 10, and 30 µg/mL DEP-treated microglial cells with 30 min imaging intervals. (**C**) Bar graphs for the integrated red channel fluorescence intensity at 2 h timepoint comparing 3, 10, and 30 µg/mL DEP to solvent medium. (**D**) Representative images of primary microglial cells stained with Lysoview 540 (red) and 488 (green) after treatment with 3, 10, and 30 µg/mL DEP for 2 h. Scale bar = 100 µm. (**E**) Bar graphs for the integrated fluorescence intensity of Lysoview 540 to 488 in microglia cells after indicated treatment. All data are mean ± SD with one-way ANOVA. \*\*\*\**p* < 0.0001. n=3 independent experiments with 3 wells per condition.

To corroborate the findings on the lysosomal defects in microglia in response to DEP exposure, we next employed dual fluorescence labeling with Lysoview 488 and Lysoview 540. While both probes accumulate in acidic organelles, Lysoview 540 provides additional sensitivity to pH changes. Two hours of exposure to DEP did not change the integrated fluorescence intensity of Lysoview 488, while the intensity of the red Lysoview 540 fluorescence was decreased in a concentration-dependent manner (**Figure 3D**). The ratio of Lysoview 540 to Lysoview 488 signal was significantly reduced in DEP-treated microglia (**Figure 3E**). These data confirm that DEP exposure leads to lysosomal alkalinization, rather than changes in lysosomal content, thereby disrupting the acidic environment essential for lysosomal function.

### 3.4. DEP exposure increases intracellular ROS production

Impaired lysosomal acidification can enhance cytokine expression and promote a pro-inflammatory state in microglia, prompting us to investigate whether DEP exposure alters intracellular ROS levels. While physiological ROS production in immune cells supports host defense, excessive ROS accumulation can induce oxidative stress, exacerbating neuroinflammation and neuronal loss in neurodegenerative disorders [70, 71]. To monitor intracellular ROS in real time, microglia were incubated with CellROX Deep Red dye, while being exposed to increasing DEP concentrations. As shown in **Figure 4A**, DEP-exposed microglia exhibited progressively increasing red fluorescence over time, and this increased intensity correlated with DEP concentration. Time-course quantification of integrated fluorescence intensity confirmed a dose-dependent increase in ROS levels (**Figure 4B**). At 24 hour, integrated fluorescence intensity analysis revealed statistically significant increase in ROS levels across all DEP concentrations (**Figure 4C**). Collectively, these findings demonstrate that DEP exposure induces sustained, concentration-dependent intracellular ROS accumulation in microglia, suggesting a mechanistic link between lysosomal dysfunction and oxidative stress.

**Figure 4.**
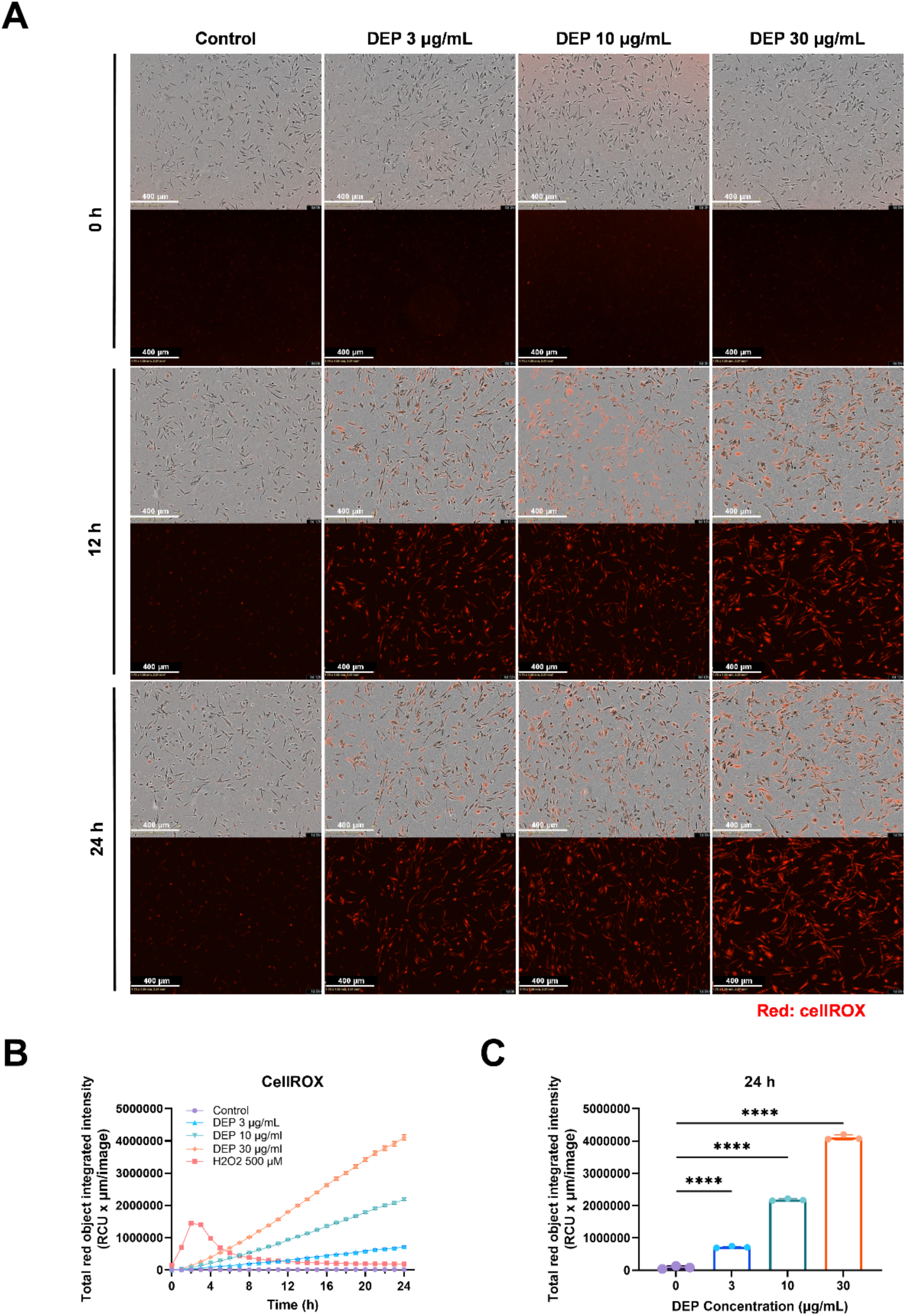
DEP exposure increases intracellular ROS production in primary mouse microglia. (**A**) Representative phase-contrast and red fluorescence images of primary microglial cells stained with CellROX Deep Red dye, treated with solvent control or 3, 10, or 30 µg/mL DEP at 0, 12, and 24 h. Scale bar = 400 µm. (**B**) Time-curves of integrated red fluorescence intensity in microglia treated with 3, 10, and 30 µg/mL DEP. (**C**) Bar graph showing integrated fluorescence intensity of CellROX Deep Red at 24 h. All data are mean ± SD with one-way ANOVA. \*\*\*\**p* < 0.0001. n=3 independent experiments with 3 wells per condition.

### 3.5. DEP exposure impairs the clearance of amyloid beta in mouse and human iMGL cells

Microglia are recognized as key players in AD pathology, attributed to their role in the clearance of amyloid beta (Aβ) aggregates [32, 48, 63]. The human iPSC-derived microglia, iMGL model is being increasingly employed to study microglial dysfunction in neurodegenerative diseases due to their translational value [32, 51, 64, 72]. To determine whether DEP exposure could interfere with the critical microglial function of clearance of Aβ, we assessed the Aβ uptake in both primary mouse microglia and human iMGL cells. Cell imaging revealed a DEP concentration-dependent decrease in green fluorescence intensity, indicating reduced Aβ uptake in both mouse (**Figure 5A and Supplementary figure S3**) and human microglial cells (**Figure 5D**), consistent with impaired Aβ clearance. Quantification of these results confirmed significant reductions in intracellular HiLyte™ Fluor 488-labeled Aβ (1-42) fluorescence intensity in mouse microglia (**Figure 5B**) and human iMGL cells (**Figure 5E**) compared to the solvent controls. Live-cell imaging demonstrated a time-dependent reduction in Aβ internalization following DEP exposure (**Figure 5C and S3**) in primary mouse microglia. Exposure of human iMGL to DEP significantly reduced their phagocytic capacity for pHrodo-labeled particles (**Figure 5F and G**). Collectively, these findings indicate that DEP exposure disrupts the microglial capacity to clear Aβ, in both mouse primary microglia and human iMGL cells, providing a potential mechanistic link between environmental particulate exposure and impaired microglial function in AD.

**Figure 5.**
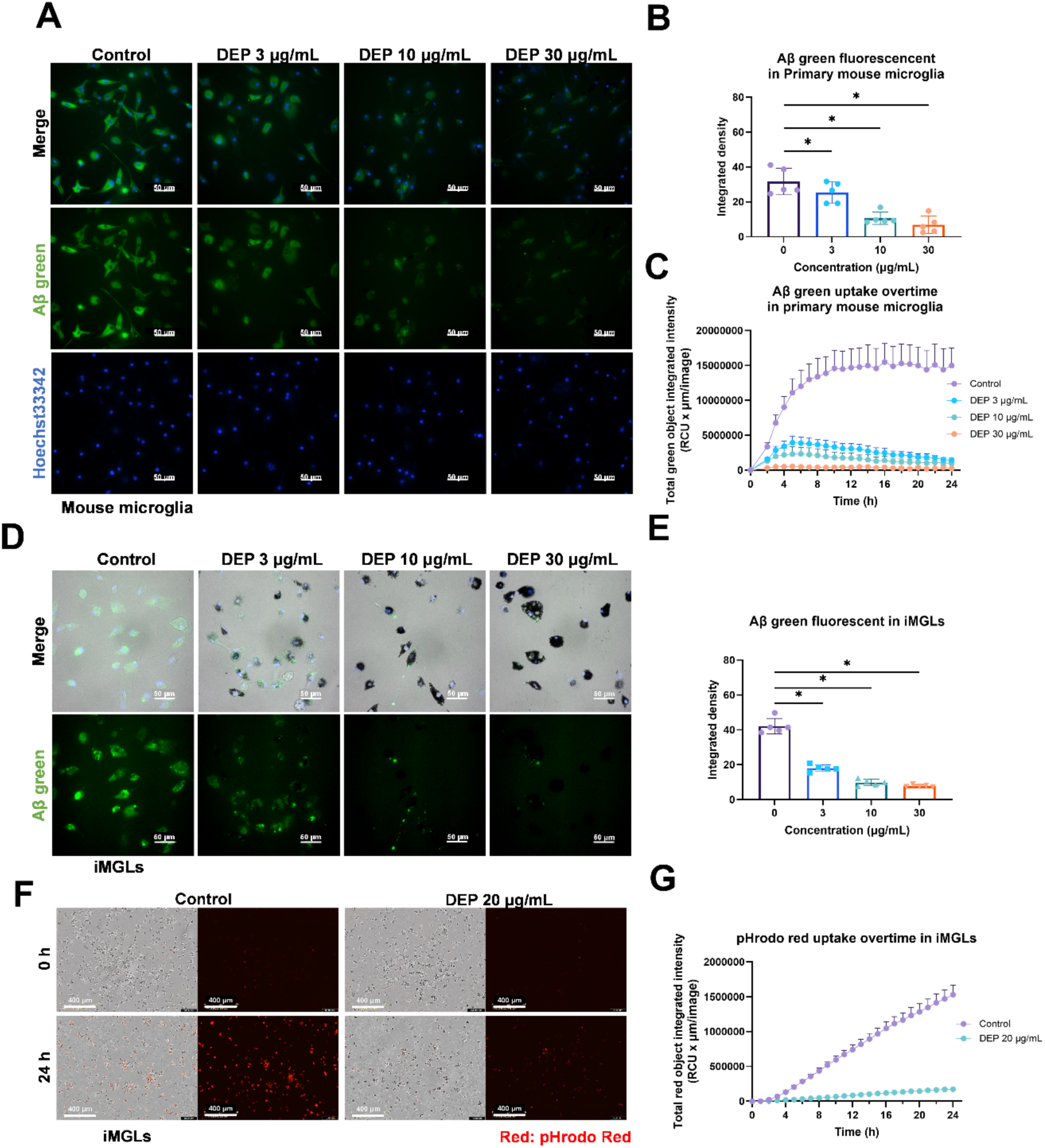
DEP exposure decreases Aβ phagocytosis in both primary mouse and human microglial cells. (**A**) Representative images of primary microglia showing HiLyte™ Fluor 488-labeled Aβ (1–42, green) uptake in cells treated with 3, 10, or 30 µg/mL DEP at 0, 12, and 24 h. Scale bars 200 µm. (**B**) Bar graphs for the integrated fluorescence intensity of HiLyte™ Fluor 488-labeled Aβ (1-42) at 24 h in mouse microglia. (**C**) Time-curves of integrated green fluorescence intensity for 3, 10, and 30 µg/mL of DEP-treated mouse microglial cells. (**D**) Representative images of iMGL cells showing HiLyte™ Fluor 488-labeled Aβ (1–42, green) uptake in cells treated with 3, 10, or 30 µg/mL DEP. (**E**) Bar graphs for the integrated fluorescence intensity of HiLyte™ Fluor 488-labeled Aβ (1-42) at 24 h in mouse microglia. (**F**) Representative images of iMGL with phase and red fluorescent channel showing phagocytosed red pHrodo red *E.coli* bioparticles with solvent medium and 20 µg/mL DEP at 0 and 24 h. (**G**) Time-curve of integrated red fluorescence intensity iMGL pretreated with 20 µg/mL DEP. All data are mean ± SD with one-way ANOVA. \**p* < 0.05 n=3 independent experiments.

### 3.6. Early transcriptional reprogramming of microglia following 4-hour DEP exposure

To dissect the early cellular responses to DEP exposure, we performed bulk RNA sequencing of mouse and human microglial cells following a 4-hour of DEP exposure. The analysis revealed profound transcriptional reprogramming with distinct pathway-specific responses in both mouse **(Figure 6A, B)** and human **(Figure 6F, G)** cells. Mouse microglia displayed a broad transcriptional response, with 571 differentially expressed genes (DEG) primarily associated with the regulation of antiviral defense pathways and a broad activation of inflammation-related genes **(Figure 6C).** There was a significant downregulation of several pro-inflammatory mediators, including tumor necrosis factor (*Tnfa*), interleukin 1 beta (*Il1b*), and interleukin 6 (*Il6*). This suppression extended to key signaling components including myeloid differentiation primary response 88 (*Myd88*), signal transducer and activator of transcription 1 and 2 (*Stat1, Stat2*), and NLR family pyrin domain containing 3 (*Nlrp3*). This suggests that DEP might have induced an immunosuppressive effect on the cells, as evidenced by downregulation of absent in melanoma 2 (*Aim2*) and Z-DNA binding protein 1 (*Zbp1*), both of which are critical components for inflammasome activation.

**Figure 6.**
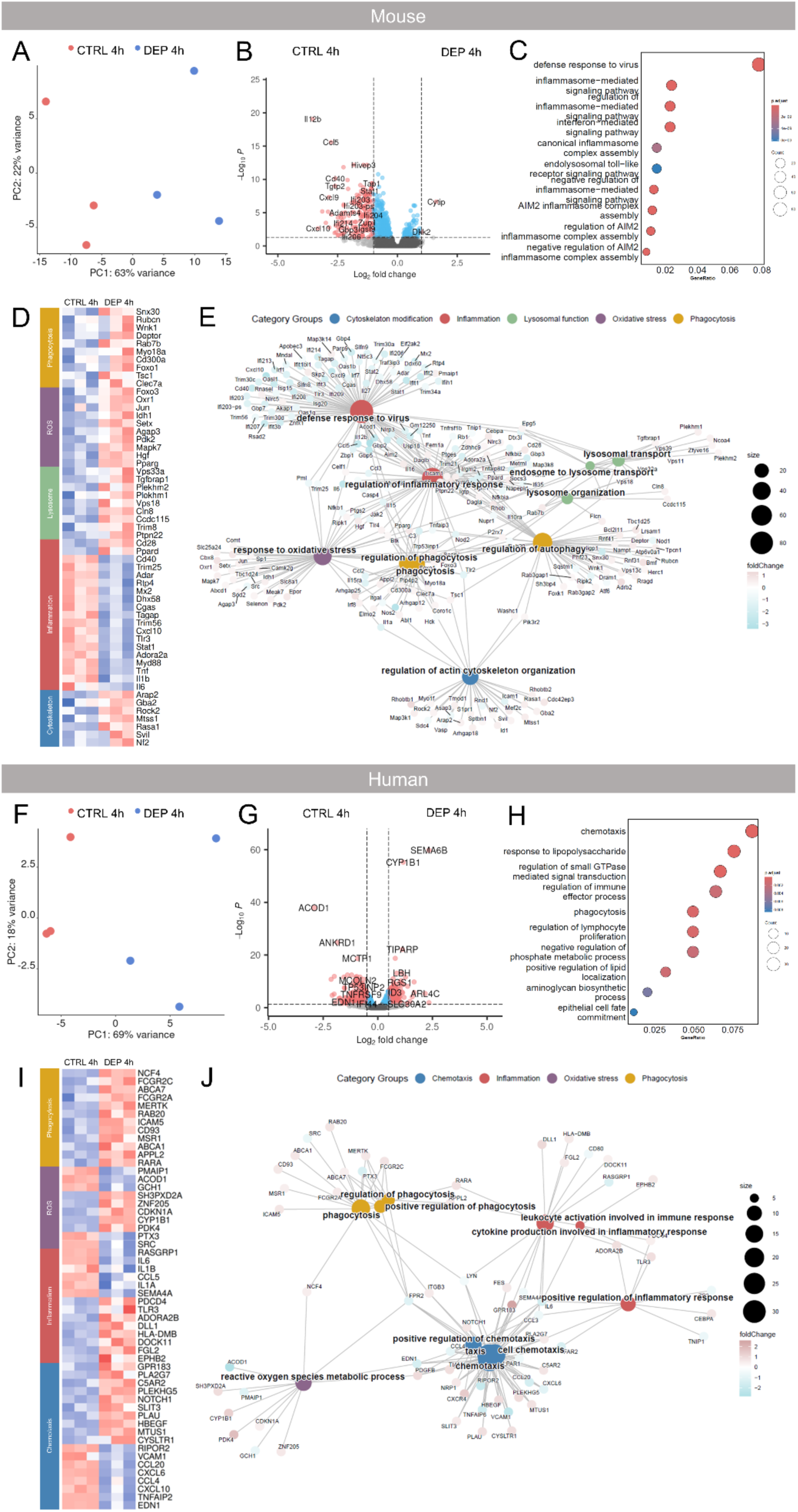
Transcriptional analysis of mouse microglial and iMGL cells following a 4-hour DEP exposure. Principal component analysis of mouse microglial **(A)** and iMGL **(F)** cells following a 4-hour DEP exposure compared to untreated cells. Volcano plots presenting fold changes and Log2 of the adjusted p-value per gene for comparing control and 4h DEP-exposed microglia in mouse **(B)** and iMGL cells **(G).** Pathway enrichment analysis showing dot plot for enriched gene clusters in mouse microglia **(C)** and iMGL **(H)** cells. Cnetplot showing selected enriched GO families, clustered by function in both mouse microglia **(D)** and iMGL **(I).** Upregulated genes are colored in pink, and downregulated genes are colored in blue. The gene clusters are Padj <0.05 with color-coded log2fold changes. Heatmaps displaying selected genes clustered by familial function for both mouse microglia **(E)** and iMGL **(J)** cells. Upregulated genes are colored in red, and downregulated genes are colored in blue. n = 3 independent experiments per condition.

Conversely, several genes involved in phagocytosis were upregulated, including forkhead box O3 (*Foxo3*), C-type lectin domain family 7-member A (*Clec7a*), TSC complex subunit 1 (*Tsc1*), forkhead box O1 (*Foxo1*), *Cd300a*, and myosin XVIIIA (*Myo18a*).

This gene upregulation was further supported by the increased expression of *Rab7b*, DEP domain containing MTOR interacting protein (*Deptor*), WNK lysine deficient protein kinase 1 (*Wnk1*), Rubicon autophagy regulator (*Rubcn*), and sorting nexin 30 (*Snx30*). Lysosomal function genes also showed upregulation, specifically protein tyrosine phosphatase non-receptor type 22 (*Ptpn22*), tripartite motif-containing 8 (*Trim8*), coiled-coil domain containing 115 (*Ccdc115*), *Cln8*, vacuolar protein sorting 18 and 33A (*Vps18, Vps33a*), pleckstrin homology and RUN domain containing M1 and M2 (*Plekhm1, Plekhm2*), and TGF-beta receptor associated protein 1 (*Tgfbrap1*). Notably, the upregulation of phagocytic and lysosomal genes stands in contrast to the functional impairment observed in **figure 2** and **figure 3**, suggesting that microglia undergo a high-effort attempt to clear DEP at the transcriptional level, but are ultimately overwhelmed by the metabolic stress and cellular exhaustion, leading to functional failure.

Similarly, stress-related genes were also upregulated, including peroxisome proliferator-activated receptor gamma (*Pparg*), Jun proto-oncogene (*Jun*), oxidation resistance 1 (*Oxr1*), hepatocyte growth factor (*Hgf*) or ROS regulators via metabolic pathways, such as pyruvate dehydrogenase kinase isozyme 2 (*Pdk2*), calcium/calmodulin-dependent protein kinase II gamma (*Camk2g*), isocitrate dehydrogenase (*Idh*), mitogen-activated protein kinase 7; ERK5 (*Mapk7*) and others (**Figure 6D, E**). This coordinated gene upregulation might indicate a compensatory effort to clear DEP particles, however, as shown in figure 2, the cells exhibit a functional impairment in their capacity to process the DEP particles. Furthermore, the upregulation of cytoskeleton genes, including neurofibromin 2 or Merlin (*Nf2*) supervillain (*Svil*), metastasis suppressor 1; also known as MIM, Missing-in-Metastasis (*Mtss1*), Rho-associated coiled-coil containing protein kinase 2 (*Rock2*), ArfGAP with RhoGAP domain, ankyrin repeat and PH domain 2 (*Arap2*) aligns with earlier observations of significant morphological changes following DEP exposure **(Figure 1).**

In iMGL cells, 438 genes were differentially expressed after 4 hours of DEP exposure. Pathway enrichment analysis showed regulation of phagocytosis, chemotaxis, and immune response-related families **(Figure 6H).** Phagocytosis-related genes expression were upregulated, including macrophage scavenger receptor 1 (*MSR1*), MER proto-oncogene tyrosine kinase (*MERTK*), Fc gamma receptor IIa (*FCGR2A*), cluster of differentiation 93 (*CD93*), adaptor protein, phosphotyrosine interaction, PH domain and leucine zipper containing 2 (*APPL2*), RAB20, ATP-binding cassette transporter A1 and A7 (*ABCA1, ABCA7*), Neutrophil cytosolic factor 4; p40phox (*NCF4*), and Fc gamma receptors IIa and IIc (*FCGR2A, FCGR2C*), suggesting a high effort to engulf the diesel particles **(Figure 6I, J).** The chemotactic response was characterized by the downregulation of several key ligands and adhesion molecules, including endothelin 1 (*EDN1*), C-C motif chemokine ligands 4 and 20 (*CCL4, CCL20),* C-X-C motif chemokine ligand 6 (*CXCL6*), vascular cell adhesion molecule 1 (*VCAM1*), and RHO family interacting cell polarization regulator 2 (*RIPOR2*). However, there was an upregulation of receptors and signalling scaffolds, such as cysteinyl leukotriene receptor 1 (*CYSLTR1*), microtubule associated scaffold protein 1 (*MTUS1*), heparin binding EGF-like growth factor (*HBEGF*), urokinase-type plasminogen activator (*PLAU*), slit guidance ligand 3 (*SLIT3*), notch receptor 1 (*NOTCH1*), pleckstrin homology and RhoGEF domain containing G5 (*PLEKHG5*), complement C5a receptor 2 (*C5AR2*), phospholipase A2 group VII (*PLA2G7*), and G protein-coupled receptor 183 (*GPR183*). This divergent expression profile may indicate a disrupted attempt by microglia to migrate toward damaged cells. Inflammatory genes such as *IL6*, semaphorin 4A (*SEMA4A*), *CD80*, and RAS guanyl releasing protein 1 (*RASGRP1*) were downregulated. In contrast, upregulated inflammatory-related genes included ephrin type-B receptor 2 (*EPHB2*), fibrinogen-like 2 (*FGL2*), dedicator of cytokinesis 11 (*DOCK11*), major histocompatibility complex class II DM beta (*HLA-DMB*), delta-like canonical Notch ligand 1 (*DLL1*), adenosine A2b receptor (*ADORA2B*), Toll-like receptor 3 (*TLR3*), and programmed cell death 4 (*PDCD4*). Similarly, oxidative stress-related genes showed a split response; with pyruvate dehydrogenase kinase 4 (*PDK4*), cytochrome P450 family 1 subfamily B member 1 (*CYP1B1*), cyclin-dependent kinase inhibitor 1A (*CDKN1A*), zinc finger protein 205 (*ZNF205*), and SH3 and PX domains 2A (*SH3PXD2A*) being upregulated, while pentraxin 3 (*PTX3*), GTP cyclohydrolase 1 (*GCH1*), aconitate decarboxylase 1 (*ACOD1*), and phorbol-12-myristate-13-acetate-induced protein 1 (*PMAIP1*) being downregulated.

Taken together, these early transcriptional changes indicate that short-term DEP could potentially shift away from a pro-inflammatory phenotype toward a transcriptionally reprogrammed state characterized by an increase in phagocytosis, ROS, lysosome and cytoskeleton-related gene expression, in mouse microglia. Similarly, in human iMGLs, this reprogrammed state is characterized by the upregulation of phagocytic-related genes, altered ROS signalling, alongside chemotactic and some inflammatory gene regulation. These findings suggest that while microglia prioritize the engulfment and processing of foreign exhaust particles, the alternations in immune pathways and the upregulation of cytoskeletal-remodelling genes reflect a state of cellular stress that may underlie the observed functional impairments in particle clearance.

### 3.7 Divergent transcriptional reprogramming of microglia after 24 hours of DEP exposure

Sustained 24-hour exposure to DEP revealed distinct transcriptional regulation between mouse microglia cells **(Figure 7A, B)** and iMGL **(Figure 7F, G)**. At 24-hour, mouse microglia response narrowed to 297 DEGs, primarily characterized by a regulation of antigen presenting families and inflammatory processes **(Figure 7C).** Pathway enrichment analysis also revealed processes related to the regulation of nuclear division, chromosome segregation, and antiviral responses.

**Figure 7.**
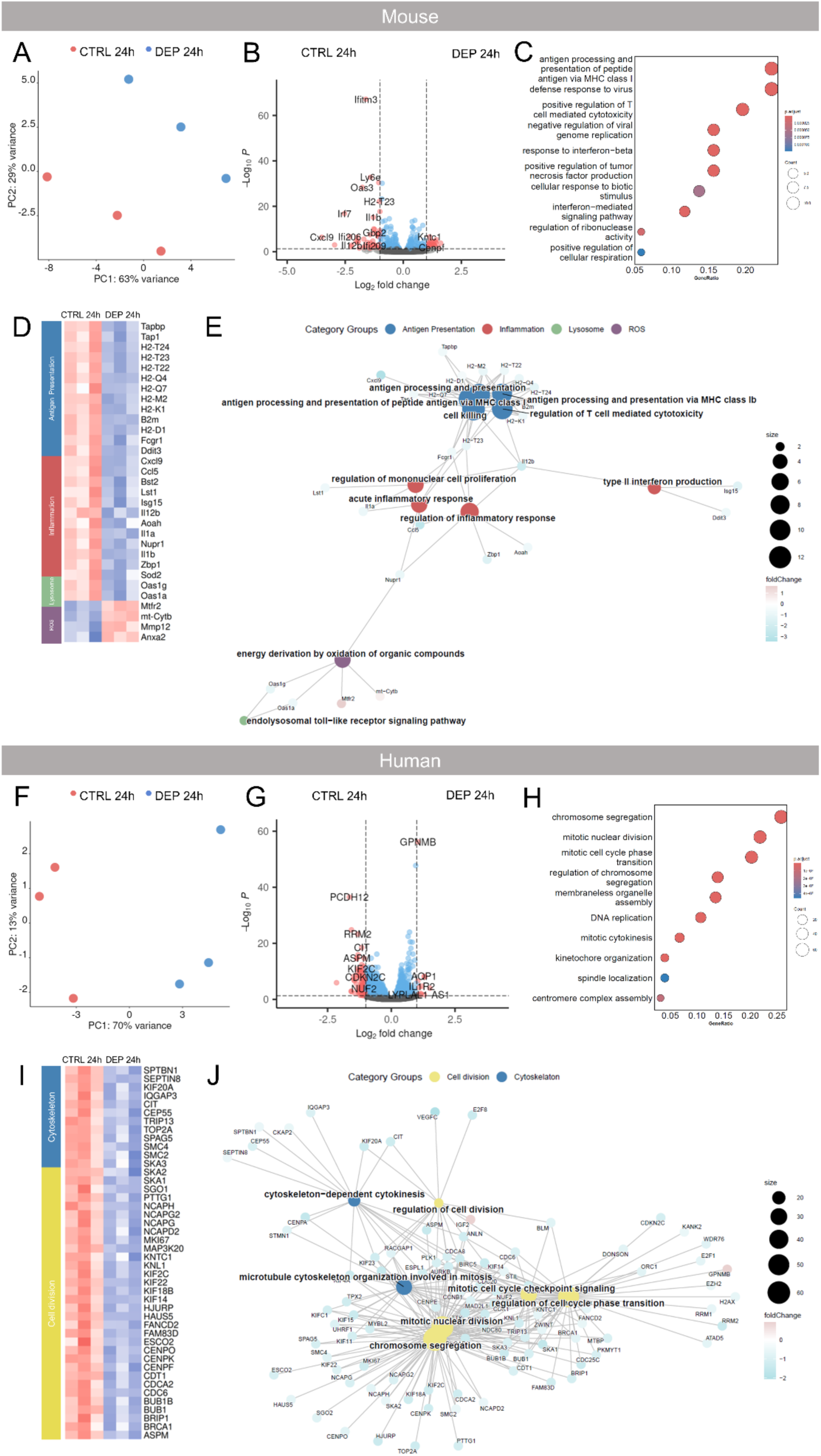
Transcriptional analysis of mouse microglial and iMGL cells following a 24-hour DEP exposure. Principal component analysis of mouse microglial **(A)** and iMGL **(F)** cells following a 24-hour DEP exposure compared to untreated cells. Volcano plots presenting fold changes and Log2 of the adjusted p-value per gene for comparing control and 24h DEP-exposed microglia in mouse **(B)** and iMGL cells **(G).** Pathway enrichment analysis showing dot plot for enriched gene clusters in mouse microglia **(C)** and iMGL **(H)** cells. Cnetplot showing selected enriched GO families, clustered by function in both mouse microglia **(D)** and iMGL **(I).** Upregulated genes are colored in pink, and downregulated genes are colored in blue. The gene clusters are Padj <0.05 with color-coded log2fold changes. Heatmaps displaying selected genes clustered by familial function for both mouse microglia **(E)** and iMGL **(J)** cells. Upregulated genes are colored in red, and downregulated genes are colored in blue. n = 3 independent experiments per condition.

Several classical inflammatory genes and antigen presenting families were downregulated. These pro-inflammatory genes include interleukin 1 alpha (*Il1a),* Z-DNA binding protein 1 (*Zbp1*), nuclear protein 1 (*Nupr1*), interferon-stimulated gene 15 (*Isg15*), the leukocyte-specific transcript 1 (*Lst1*), bone marrow stromal cell antigen 2 (*Bst2*), C-C motif chemokine ligand 5, encoding for RANTES (*Ccl5*), and C-X-C motif chemokine ligand 9 *(Cxcl9*) suggesting a failure of immune surveillance in the DEP-treated cells.

Consistently, there was a significant decrease in transcripts associated with antigen processing and inflammation-related genes, such as Fc gamma receptor 1 (*Fcgr1*), DNA Damage Inducible Transcript 3 (*Ddit3*) encoding for key transcription factor CHOP or GADD153, were also downregulated. This suppression was further reflected in a broad downregulation of major histocompatibility complex (MHC) class I family members, including histocompatibility 2 (H2-K1, H2-D1, H2-T23, H2-T22, H2-Q7, H2-Q4, and H2-M2) and transporter associated with antigen processing 1 (*Tap1*), and TAP binding protein (*Tapbp*). The interferon-induced oligoadenylate synthetase (OAS) family (*Oas1a* and *Oas1g*) genes are also downregulated. This suppression suggests a potential failure of immune surveillance in microglia following prolonged DEP exposure.

In contrast, the significant upregulation of mitochondrial and oxidative stress-related genes, specifically mitochondrially encoded cytochrome b (mt-Cytb), mitochondrial fission regulator 2 (Mtfr2), correspond to the ROS increase observed in Figure 4, indicating that the challenged cells could undergo severe metabolic strain and mitochondrial remodelling **(Figure 7D, E).**

In addition, genes related to extracellular matrix remodelling, such as matrix metalloprotein 12 (*Mmp12*) were upregulated in response to DEP challenge. At the same time, DEP also increased the expression of genes involved in cytoskeletal remodelling such as annexin A2 (*Anxa2*). Together, these findings suggest that mouse microglia shift towards a state of suppressed pro-inflammatory signalling but participate in chemotaxis, phagocytosis, and structural remodelling.

Specifically, members of the kinesin family, including kinesin family members *KIF14, KIF18A, KIF18B, KIF22, KIF2C*, and the non-SMC condensin I/II complex subunits *NCAPD2, NCAPG, NCAPG2,* and *NCAPH* were suppressed. This was accompanied by the downregulation of spindle and kinetochore associated proteins *SKA1, SKA2*, and *SKA3*, as well as centromere proteins *CENPF, CENPK,* and *CENPO*, and the mitotic checkpoint kinase *BUB1B*, pointing towards the profound dismantling effect of DEP on the mitotic machinery of the cells. Furthermore, cytoskeleton genes thyroid hormone receptor interactor 13 (*TRIP13*), centrosomal protein 55 (*CEP55*), citron rho-interacting serine/threonine kinase (*CIT*), cytoskeleton associated protein 2 (*CKAP2*), IQ motif containing GTPase activating protein 3 (*IQGAP3*), kinesin family member 20A (*KIF20A*), septin 8 (*SEPTIN8*), and spectrin beta, non-erythrocytic 1 (*SPTBN1*) were also significantly downregulated suggesting loss of essential structural scaffolds, preventing microglia from migrating **(Figure 7I, J)**. Consistent with the structural alternations, the cells transitioned into a pathological state indicated by the significant upregulation of glycoprotein Nmb (*GPNMB*) **(Figure 7G),** as well as *ANXA2* and *MMP12* **(Supplementary figure 4A),** key markers of Disease-Associated Microglia (DAM). This was further supported by an oxidative stress response, evidenced by the induction of superoxide dismutase 2 (*SOD2*), heme oxygenase 1 (*HMOX1*), and neutrophil cytosolic factors (*NCF1, NCF2, NCF4*). Inflammatory genes *CCL2*, *CCR1*, interleukin 10 (*IL10*), and SLAM family member 7 (*SLAMF7*), while the homeostatic marker C-X3-C motif chemokine receptor 1 (*CX3CR1*) was decreased. Furthermore, the cells upregulated several genes associated with autophagy and lysosomal function, including microtubule associated protein 1 light chain 3 beta (*MAP1LC3B*), sequestosome 1 (*SQSTM1*), *LAMP1*, cathepsins B and D (*CTSB, CTSD*), *RAB7B*, and *CD63* **(Supplementary figure 4A).** The upregulation of these phagocytic genes differs to the previous results showed in **figure 5F and G**, indicating a high-effort transcriptional response that does not translate into effective particle clearance.

### 3.8 DEP exposure disrupts microglial gene programs involved in amyloid-â clearance and increases inflammatory gene signature

To mimic the interaction between environmental DEP exposure and Aβ pathology, we challenged DEP-primed microglia with Aβ. We aimed to investigate whether environmental risk factors exacerbate or alter microglial gene programs associated with inflammation, clearance, and neurodegeneration. Sequential exposure of mouse microglia to DEP for 24 hours, followed by a 4-hour challenge with Aβ, induced distinct transcriptional profiles in both mouse **(Figure 8A, B)** and human iMGL **(Figure 8F, G)** cells.

**Figure 8.**
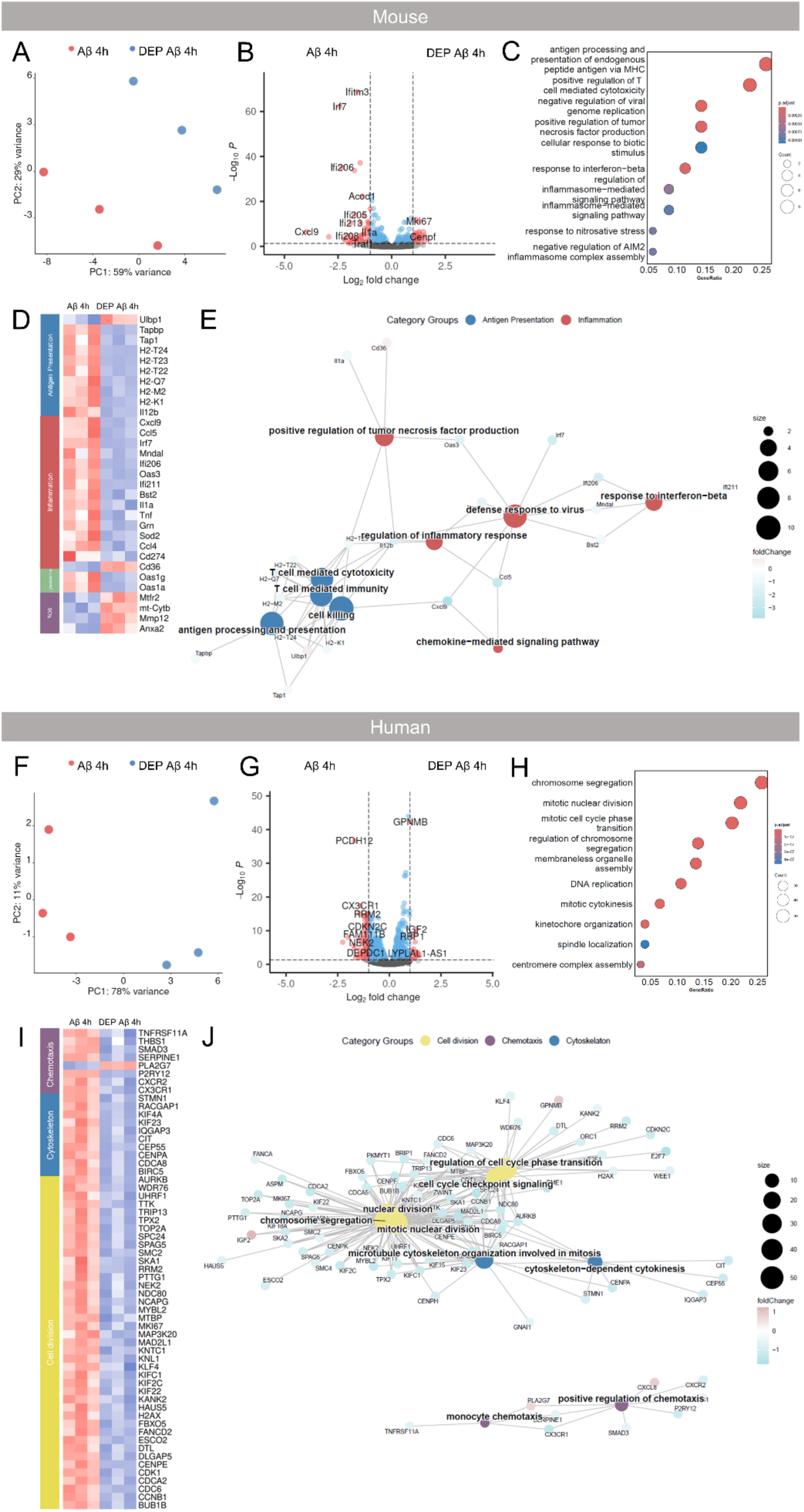
Transcriptional analysis of mouse microglial and iMGL cells following a 24-hour DEP exposure followed by 4 h of Aβ (hereby mentioned as DEP Aβ 4h condition). Principal component analysis of mouse microglial (A) and iMGL (F) cells following a 4-hour Aβ exposure compared to untreated cells. Volcano plots presenting fold changes and Log2 of the adjusted p-value per gene for comparing 4h Aβ and 24h following a 4h Aβ-exposed microglia in mouse (B) and iMGL cells (G). Pathway enrichment analysis showing dot plot for enriched gene clusters in mouse microglia (C) and iMGL (H) cells. Cnetplot showing selected enriched GO families, clustered by function in both mouse microglia (D) and iMGL (I). Upregulated genes are colored in pink, and downregulated genes are colored in blue. The gene clusters are Padj <0.05 with color-coded log2fold changes. Heatmaps displaying selected genes clustered by familial function for both mouse microglia (E) and iMGL (J) cells. Upregulated genes are colored in red, and downregulated genes are colored in blue. n = 3 independent experiments per condition.

In mouse microglia, 173 DEGs were identified, with transcriptional changes primarily associated with antiviral signalling and inflammatory pathways **(Figure 8C)**. Similar to 24-hour DEP exposure alone, we observed a significant downregulation of antigen presenting genes *Tap1, Tapbp,* and MHC class I family (*H2-K1, H2-M2, H2-Q7, H2-T23, H2-T22, H2-T24)*. While scavenger receptor such as Collectin sub-family member 12 (*Colec12*) and *CD36* was upregulated, the majority of inflammatory genes *Il1a,* Tetherin (*Bst2*), interferon-activated gene 211 (*Ifi211*), *Ifi206, Oas3,* myeloid nuclear differentiation antigen-like (*Mndal*), interferon regulatory actor 7 (*Irf7*), *Ccl5* and *Cxcl9* were significantly downregulated, suggesting an altered immune response, and adaptive changes in alternate cellular clearance routes **(Figure 8D, E)**. Overall, this might indicate a reduced degradation and clearance capacity and an altered immune response.

DEP-exposed iMGLs followed by Aβ challenge resulted in a profound transcriptional shift, with 1,176 identified DEGs. Pathway enrichment analysis revealed a significant downregulation of gene families critical for nuclear division, chromosome segregation, and cell division families **(Figure 8H).**

iMGL challenged by Aβ showed a similar decrease in gene expression, with KIF family (*KIF2C, KIF22, KIFC1*), centromere related genes (*CENPE*, Kinetochore Null 1 (*KNL1*), *NDC80, SPC24,* kinetochore associated 1 (*KNTC*), *BUB1B*), cell cycle regulators (such as, cyclin dependent kinase 1 (*CDK1*), cell division cycle 6 (*CDC6*), cyclin B1 (*CCNB1*), cell division cycle associated 2 (*CDCA2*)) and others, being downregulated, suggesting continuous failure of cell division. Similarly, cytoskeleton-related genes and chromosome segregation-related genes were downregulated, including kinesin-like protein 23 (*KIF23*), Rac GTPase-activating protein 1 (*RACGAP1*), and Stathmin, structural microtubule-associated protein 1 (*STMN1*), aurora kinase B (*AURKB*), Survivin, baculoviral inhibitor of apoptosis repeat-containing 5 (*BIRC5*), centromere protein A (*CENPA*), centrosomal protein of 55 kDa (*CEP55*), citron Rho-interacting serine/threonine kinase (*CIT*), IQGAP3, pointing on a potential dysfunctional migration mechanism and a significant impairment in structural remodelling. This was accompanied by a collapse of the chemotactic recruitment, immune cell migration, adhesion-associated genes, evidenced by the reduced expression of *CX3CR1*, *CXCR2*, Purinergic Receptor P2Y12 (*P2RY12*), Serpin Family E Member 1 (*SERPINE*), SMAD Family Member 3 (*SMAD3*), Thrombospondin 1 (*THBS1*), and *TNFRSF11A* **(Figure 8I, J).**

Interestingly, iMGL exhibited an upregulation of genes associated with oxidative stress and ferroptosis, including Glutathione Peroxidase 4 (*GPX4*), Nuclear Factor Erythroid 2-Like 2 (*Nrf2*), *HMOX1*, *SOD2*, and *PLA2G7*. This was accompanied with the activation of lysosomal and autophagic genes Cathepsins B, D, and K (*CTSB, CTSD, CTSK*), *SQSTM1*, Alpha-L-Fucosidase 1 (*FUCA1*), Progranulin (*GRN*), *CD63*, and *GPNMB*. Furthermore, the cells exhibited a pro-inflammatory profile with a significant increase of *TNF*, *CXCL8/IL-8*, *CCL4*, *IL10*, *CD274*, TNF Alpha Induced Protein 2 (*TNFAIP2*), and Galectin 3 (*LGALS3*) genes. Membrane homeostasis markers Apolipoprotein E (*APOE*) was also upregulated, while Bridging Integrator 1 (*BIN1*) was uniquely downregulated, suggesting a reorganization of lipid and membrane dynamics in response to the combined DEP and Aβ challenge **(Supplementary figure 4B).** This shift at the level of transcription suggests an attempt of the iMGL to engulf the Aβ particles, however, as shown earlier in **Figure 5 D** and **E**, the cells fail to internalize and clear Aβ, that could lead to an accelerated amyloid pathology due to a reduction in microglial capacity for Aβ clearance and increased vulnerability to neurodegeneration.

### 3.9 DEP exposure reprograms microglial metabolic pathways over time

To further elucidate how DEP exposure could alter microglial energy metabolism over time and impact their functional capacity to respond to pathological stressors such as Aβ, we assessed the potential changes in metabolic pathways, such as glycolysis, tricarboxylic acid cycle (TCA), and fatty acid oxidation (**Supplementary figures S5, S6, and S7**).

Bulk RNA sequencing revealed that the metabolic transcriptome of mouse microglia is altered during the DEP exposure (**Supplementary figure S5**). Hexokinase 2 (*Hk2*), a key enzyme for glucose phosphorylation, is downregulated following 4-hour DEP exposure, but reaches the baseline level at 24-hour (**Figure 9**). 6-phosphofructo-2-kinase/fructose-2,6-bisphosphatase 3 (*Pfkfb3*) was downregulated at 4-hour exposure, but restored to baseline at 24-hour. *Pfkfb3* is a vital glycolysis regulator that is an indicator of glycolytic activity. These results indicate that glycolysis is decreased at short exposure with DEP and similar to the non-challenged cells at later time points of DEP exposure. The glycolysis and mitochondrial OXPHOS link, Pyruvate dehydrogenase kinase 1 (*Pdk1*), was found upregulated at 24-hour, following the combined exposure of Aβ, favoring glycolysis over mitochondrial oxidation and limiting pyruvate entry into the TCA cycle. Parallelly, *Ldha* was upregulated at 24-hour of DEP exposure in mouse microglia. In the TCA cycle, Isocitrate dehydrogenase 1 (*Idh1*) was upregulated after Aβ exposure. Overall, mouse microglia displayed an initial inhibition of the glycolytic flux, which later shifted towards increased glycolysis due to *Pdk1*-driven pyruvate shunting and potential TCA cycle activity during DEP and Aβ exposure (**Supplementary figure S5-S7**).

**Figure 9.**
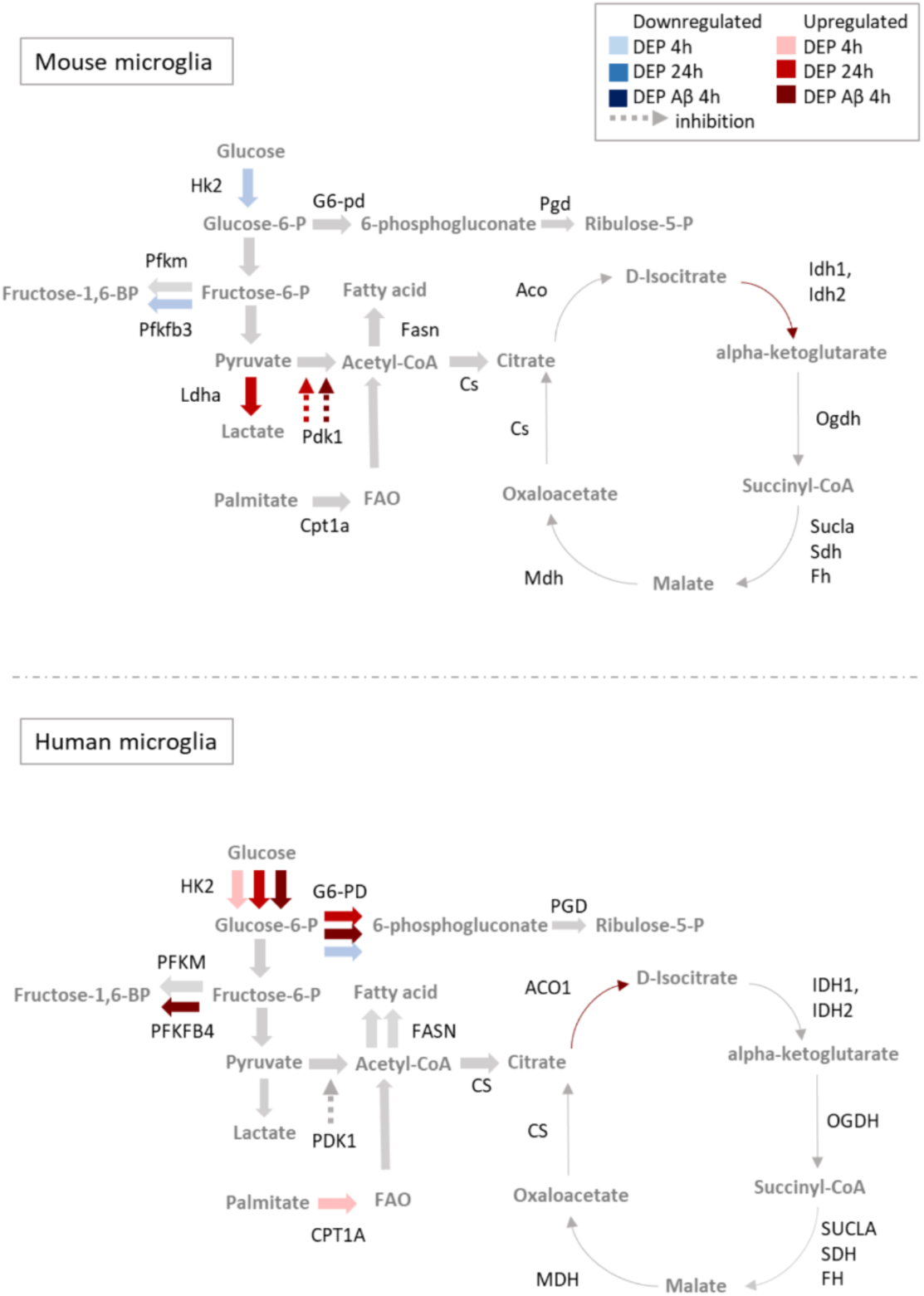
Metabolic differences and similarities in mouse and iMGL cells for glycolysis and the TCA cycle. Upregulated genes across all conditions (DEP 4-hour (depicted as DEP 4h), DEP 24-hour (depicted as DEP 24h), and DEP and Aβ 4h (depicted as DEP Aβ 4h)) are shown in a color gradient of blue, and downregulated genes across all conditions are shown as color gradients of red. Grey arrows depict unchanged pathways. The dotted arrow depicts an inhibitory pathway.

Human iMGLs responded to DEP with sustained upregulation of multiple metabolic pathways (**Supplementary figure S6**). At all measured timepoints (4-hour DEP, 24-hour DEP, and 4-hour DEP and Aβ), *HK2* was upregulated, indicating persistent initiation of glycolysis across conditions. Glucose-6-phosphate dehydrogenase (*G6-PD*), the rate-limiting enzyme of the pentose phosphate pathway (PPP) was found to be upregulated during later timepoints of 24-hour DEP and DEP followed by Aβ exposure (**Figure 9** and **Supplementary figure S7**). The increase in gene expression related to the PPP pathway indicates enhanced glucose oxidation and elevated NADPH production to potentially maintain cellular redox balance and defence against oxidative stress.

These results align with our previous research, where proteomic analysis on PPP enzymes following LPS stimulation led to an increase in Phosphogluconate dehydrogenase (PGD), an enzyme responsible for conversion to Ribulose-5-P [51], suggesting that glucose is shunted towards PPP in iMGLs cells. Fatty acid metabolism was also altered, with the key β-oxidation enzyme, Carnitine palmitoyltransferase 1A (*CPT1A),* being upregulated following 4-hour DEP exposure. Within the TCA cycle, Aconitase 1 (*ACO1*) was found to be upregulated when DEP exposure was followed by Aβ challenge, potentially inducing a metabolic shift and augmenting the DAM state of microglia (**Figure 9** and **Supplementary figure S7)**. Altogether, DEP exposure in iMGLs led to an upregulation of key metabolic pathways, including glycolysis, PPP, and fatty acid oxidation, with early responses dominated by glycolysis and fatty acid oxidation, and later responses related to additional PPP, metabolic and antioxidant programs.

## 4. Discussion

In this study, we investigated the effects of DEP exposure on transcriptome profile and microglial functions, including phenotype alterations, phagocytic capacity, lysosomal activity, ROS generation, and Aβ clearance in both mouse and human microglia. Our data showed that exposure to DEP indeed could affect both the response and key functions of microglia, thereby impairing the degradation of Aβ. Exposure to Aβ in DEP-challenged human microglia upregulated DAM-related genes and increased inflammatory and oxidative stress-related genes, coupled with a loss of homeostatic and clearance capacity. The transcriptomic responses indicated a compromised resilience in both mouse microglia and human iMGL cells upon environmental exposure in an AD-relevant setting.

DEP exposure showed that microglial cells became more ameboid, accompanied by a significant increase in the cell roundness. In single-cell analysis, microglia displayed a decrease in their length and an increase in their circularity after DEP exposure. In line with previous research, DEP-treated microglia were qualitatively larger, more spherical, and lacked processes compared to control microglia [73], although microglia observed in isolation (e.g., *in vitro* cultures) lack the intricate morphological complexity and functional states seen *in vivo* [36, 74]. In our study, DEP exposure decreased the phagocytic capacity of microglial cells in a concentration-dependent manner. Following DEP exposure, microglia engulfed DEP, potentially leading to a saturation of their phagocytic capacity and resulting in a decreased ability to phagocyte pHrodo red *E.coli* bioparticles. In accordance with these findings, a recent study demonstrated that iMGLs showed diminished phagocytic activity in response to traffic-related PM exposure [57]. EEA1 and LAMP1, markers for active endosomal and lysosomal compartments, enable precise delineation of the phagocytic cargo progression. Early endosomes, defined by EEA1, represent a critical sorting hub for internalized cargo, directing it toward recycling pathways to the plasma membrane, retrograde transport to the trans-Golgi network, or degradation in lysosomes [75]. Exposure to DEP resulted in a dose-dependent decrease in LAMP1 and EEA1 signal intensity, indicative of alterations in the early stages of the endocytic pathway. This reduction in signal intensity is consistent with impaired vesicular trafficking, lysosomal membrane permeabilization, and eventual collapse of microglial phagocytic capacity, effects paralleled by the transcriptomic profile.

Recent evidence showed that exposure to PM could induce oxidative stress in various cell types [57, 76, 77]. Acute exposure of microglial cells to high-dose PM2.5 increased mRNA expression of pro-inflammatory cytokines like IL-1β and TNF-α [78] and decreased cell survival due to neuroinflammation and ROS production [77]. Moreover, relevant to our air pollution model, the link between microglial activation and neuroinflammation was also demonstrated using PM *in vitro*. Studies demonstrated that DEP can induce microglial morphological changes, and increase ROS production, [66, 73, 79]. This increase in ROS production observed when microglial cells are exposed to DEP is in line with our hypothesis that DEP may play a role in the development and progression of neurodegenerative diseases. In the neurodegeneration context, using human *in vitro* 3D microfluidic model that incorporates neurons, astrocytes and microglia, PM has been shown to penetrate the *in vitro* BBB [25]. Conditioned medium from PM-treated neurons and astrocyte co-cultures induced microglial migration, and longer exposures increased their cytokine production, while single-cultured microglia showed a lack of immune response upon direct PM exposure, similarly to our observations. This microglial transition induced by IL-1*β* and IFN-*γ* from co-cultured neurons and astrocytes under PM_2.5_ exposure raises the concern of testing DEP in a broader neuroinflammation model, including astrocytes, to better account for its translatability. The lack of a prominent immune response upon DEP exposure in single microglial cultures aligns with this study. The degradation of Aβ by microglia was impaired in both mouse and iMGL cells following DEP exposure. Interestingly, the exposure to AgNP (silver nanoparticles) inhibited Aβ uptake by BV-2 cells [82]. The reduced ability of microglia to internalize Aβ fibrils and pHrodo red *E.coli* bioparticles suggests an impairment in microglia phagocytic function. In a recent study, DEP extracted from a modern diesel vehicle using a renewable diesel caused no difference in gene expression level and only minor functional changes in iMGLs, highlighting the importance of modern emission control measures and their impact on microglia and brain health [80]. Exposure to DEP derived from a heavy-duty diesel engine run with a fossil or renewable diesel led to immunosuppression of cytokine secretion and lysosomal defects in iMGLs, which is in line with our functional and transcriptomic results. Different types of pollutants have been reported to impair phago-lysosomal machinery in monocytic cell lines, T-cells [81–83]. Airborne particulate matter (APM) has been shown to accumulate in lysosomes, which disturbs the lysosomal pH balance in pulmonary diseases [84]. APM hinders autophagosome production and blocks autophagic flux, which remains consistent with our results.

Short DEP 4-hour exposure induced a change in microglial transcription toward a non-canonical activation state in both mouse microglia and human iMGLs. This change was characterized by a suppression of classical inflammatory pathways, and a concomitant robust induction of phagocytic, lysosomal, oxidative stress, and cytoskeletal genes. The downregulation of key pro-inflammatory mediators and inflammasome components (Tnfa/IL6/IL1b, MYD88, STAT1/2, NLRP3, AIM2, ZBP1) suggests an early immunosuppressive phenotype, at the same time as microglia is being engaged in particle engulfment and remodeling their morphology. Our data shows that short DEP exposure strongly upregulated genes involved in phagocytosis and lysosomal trafficking (Clec7a, MSR1, MERTK, FCGR2A/C, RAB20, RAB7B, Vps18/33a, Plekhm1/2, Tgfbrap1), consistent with an increase in gene expression related to internalization and processing of DEP particles. Similar increase of lysosomal and autophagy-related pathways and decrease in cytokine gene expression have been observed in microglia [80], monocytic cell lines [85], and macrophages [81] exposed to various particulate air pollutants.

Distinct oxidative metabolism regulation upon exposure to inflammatory stimuli has been observed in iMGLs [51]. Recent work in human iPSC-derived microglia exposed to traffic-related diesel particles showed induction of Toll-like receptor and oxidative stress pathways, together with impaired lysosomal function and defective waste clearance, reflecting the discrepancy between the effort of a microglial transcriptional “activation” and the functional phagocytic impairment demonstrated by our findings. This decoupling or divergence between gene-level upregulation of phagocytic machinery and reduced effective cargo clearance could lead to inefficient particle or aggregate removal, as observed by impaired Aβ uptake in DEP-exposed microglia.

The observed induction of oxidative stress and metabolism-related genes (Pparg, Jun, Oxr1, Pdk2, Camk2g, Idh, Mapk7, and multiple ROS-handling enzymes) aligns well with findings that DEP and other fine particulate pollutants could drive NADPH oxidase-dependent ROS production, mitochondrial dysfunction, and antioxidant responses in microglia and brain endothelial cells [86]. Nanometer-scale diesel particles could increase the activity of microglial NADPH oxidase, leading to ROS generation that in turn could result in dopaminergic neurotoxicity. Such fast changes in metabolic and redox state could “prime” microglia, making them more reactive but also more prone to functional collapse when faced with additional challenges, such as amyloid-β. This is consistent with our observations of impaired Aβ uptake despite a transcriptional program favouring phagocytosis and lysosome/autophagy activation.

Furthermore, the upregulation of cytoskeleton-and motility-related genes (including Nf2, Rock2, Mtss1, Arap2, and multiple kinesins at later time point) together with suppression of chemokine and adhesion ligands in iMGLs (for example, CCL4, CXCL6, VCAM1) suggests that DEP exposure could mediate extensive structural remodelling without effective chemotactic guidance, a phenomenon that could lead to impaired microglial migration around Aβ deposits in cells exposed to DEP. Indeed, following PM_2.5_ exposure in AD mice, the plaque deposition and phosphorylated tau levels in the hippocampus and cortex were increased, which prominently emphasizes the exacerbation of AD pathology due to PM_2.5_ [87]. Collectively, our data support a model in which short-term DEP exposure result in a decrease of canonical inflammatory pathways, and an increased transcription of phagocytosis-related genes. Although our data also revealed species-specific differences, short DEP exposure in mouse microglia suppressed inflammatory genes while upregulating phagocytic, lysosomal, and cytoskeletal genes, whereas in iMGLs DEP exposure induced chemotactic priming, altered ROS signalling, reduced classical inflammatory mediators, and increased phagocytosis, all these changes together may lead to defective clearance of toxic protein aggregates and increase the risk of neurodegeneration.

Both mouse and human microglia 24-hour DEP exposure result in a switch from an early compensatory response to a state of structural and functional exhaustion. While the cells remain under severe metabolic strain and increased oxidative stress, they suffer a profound decrease of core homeostatic gene expression, failure of immune surveillance and antigen presentation in mouse cells, and a decrease of the mitotic and cytoskeletal-related genes in human iMGLs. This suggests that prolonged DEP exposure induces a form of cellular block, where the microglia are metabolically unable to maintain their protective roles or successfully resolve the particle load. An additional Aβ challenge in mouse microglia resulted in robust suppression of pro-inflammatory genes and phagosome pathway genes, and an upregulation of scavenger-related genes, suggesting an alternative stress response compared to iMGL cells. These findings can explain the functional deficits caused by phagocytic defects, lysosomal alkalization, which leads to ROS accumulation in our functional results.

Moreover, Aβ challenge in DEP-exposed microglia led to an upregulation of DAM markers, inflammatory, oxidative, and antioxidative responses, as well as lysosomal and autophagy genes, indicating a highly activated state, with a potentially impaired Aβ clearance. This could indicate two different approaches of microglia towards Aβ clearance in mouse and human. Recent characterization of Aβ-associated microglia in human AD brain tissue has reported enrichment in genes for cell migration (*ITGAX, GPNMB*), phagocytosis *(COLEC12, MSR1),* lipid localization *(SPP1, PPARG)* [88]. These results support our findings on exacerbation of AD pathology due to DEP exposure in iMGLs. Transcriptionally, our results show a more pronounced effect of DEP exposure on iMGLs than on primary mouse microglia, which is consistent with the results in previous studies where DEP exposure has had more pronounced effects on iMGLs than on primary mouse microglia [80]. This could highlight the importance of studying DEP exposure effects in human-derived models due to their higher sensitivity compared to mouse microglia.

Recent studies show that mouse microglia engage in metabolic reprogramming [89, 90] and transcriptionally upregulate hexokinases early during pro-inflammatory LPS challenge, favoring glycolysis, while iMGL cells preferentially increase phosphofructokinases, suggesting species differences in glycolytic engagement during immune activation [51]. The observed early upregulation of glycolytic genes (e.g., *Hk2*, *Cpt1a*) and later activation of autophagy and oxidative stress pathways in iMGL cells exposed to DEP strongly indicates their complex immune response and engagement of metabolic and clearance pathways upon inflammatory insult. In activated microglia, glycolysis, PPP pathway, and lipid catabolism are key for rapid ATP and NADPH production, supporting immune functions such as ROS generation and cytokine synthesis [89, 90]. The upregulation of PPP genes (e.g., *G6PD*) late after DEP exposure or in Aβ challenged iMGLs, was consistent with PPP’s central role in antioxidant defense and microglial adaptation during prolonged or secondary insult. In mouse microglia, the delayed glycolytic shift (*Pdk1* upregulation at 24 h of DEP) and persistent downregulation of inflammatory genes after DEP exposure further highlight species-distinct dynamics in metabolic adaptation and inflammation.

Our results indicate that certain pathways, like lipid catabolism, autophagy, PPP, and glycolysis, are more elaborate and complex in iMGL cells than in mouse microglia. In iMGL cells, we observed an abrupt rise in anti-inflammatory and anti-oxidative genes at 24-hour DEP exposure or in combined DEP and Aβ challenge, while clearance pathways such as autophagy and scavenging stayed robustly active. Parallelly, we observed glycolysis initiation and PPP pathway activation at 4-hour and 24-hour DEP exposure, respectively. These findings collectively indicate that DEP may play a role in the pathogenesis of neurodegeneration, potentially through mechanisms involving microglial activation, increased ROS generation, autophagy, metabolism, and impairment of endo-lysosomal trafficking and degradation processes. The transcriptomic responses thus suggest an increase in the neuroinflammatory risk and a compromised resilience in iMGL cells upon environmental exposure in an AD relevant model.

## 5. Conclusion

Our research demonstrated that primary mouse microglia and human iMGLs display robust functional responses to DEP exposure, such as microglial morphology, phagocytic capacity, lysosomal activity, and ROS generation. These changes suggest that DEP exposure may have a significant impact on the homeostatic functions of microglia, thereby leading to neuroinflammation and subsequent broader neurological consequences, highlighting the potential for air pollution, especially of the traffic-related particulate matter, to exacerbate or even initiate neurodegenerative processes through sustained oxidative mechanisms and impaired Aβ aggregate uptake.

## Acknowledgements

We thank Prof. Bart J.L. Eggen for providing the human iPSC cells used in this study. AMD is the recipient of an Alzheimer Nederland grant, and a Rosalind Franklin Fellowship co-funded by the European Union and the University of Groningen.

