## Supplemental figures for "Diesel exhaust particles disrupt mouse and human iPSC-derived microglial function and Amyloid-β clearance in Alzheimer’s disease models": Supplementary figure.docx

**Supplementary Figures**

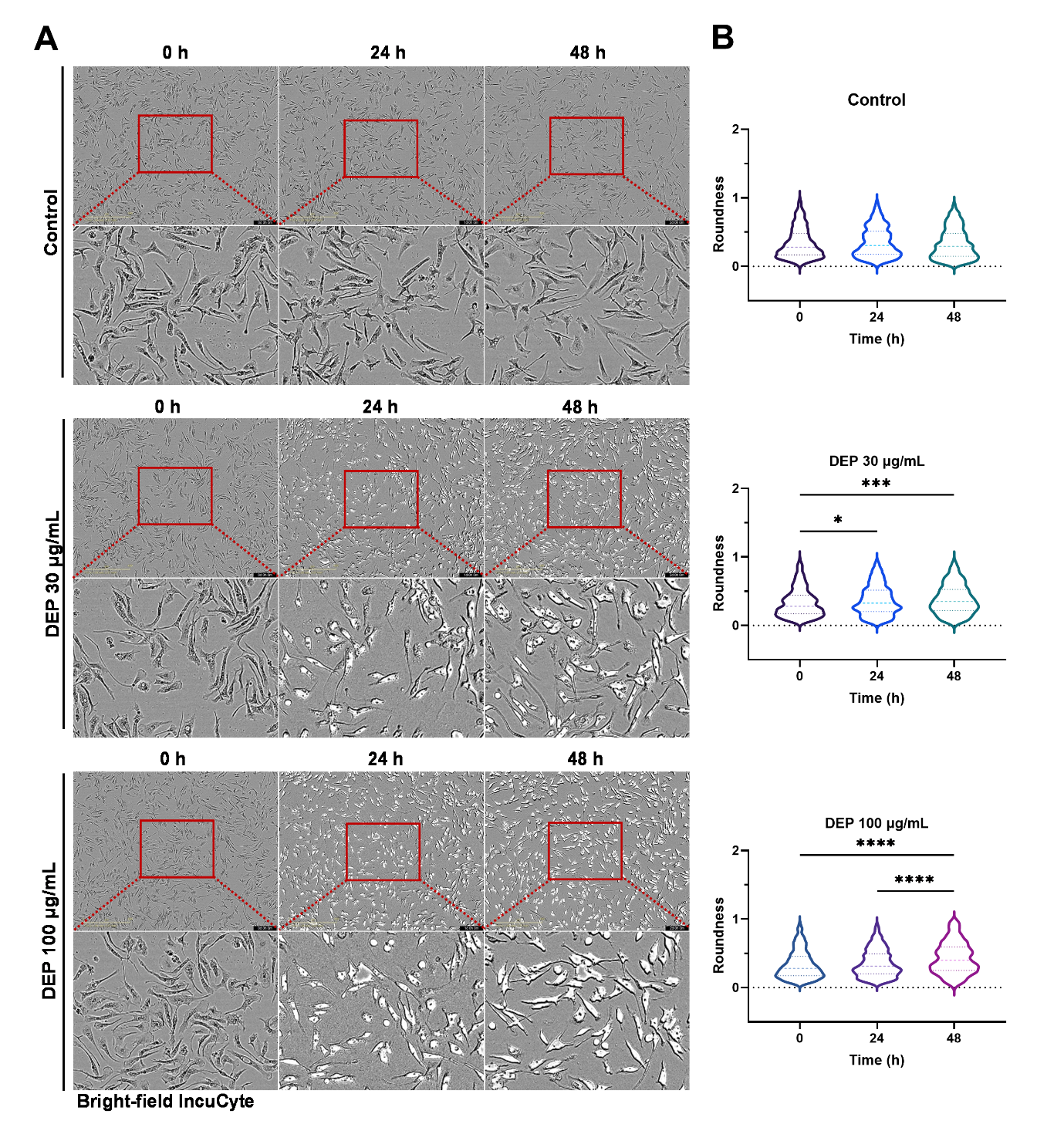

**Supplementary figure S1**. Morphology changes of primary microglial cells upon DEP exposure. (**A**) Representative phase contrast images of primary microglial cells exposed to solvent medium, 30 and 100 μg/mL DEP for 0, 24, and 48 h. Scale bar 400 μm. (**B**) Roundness quantification of microglial cells upon exposure to solvent medium or DEP.

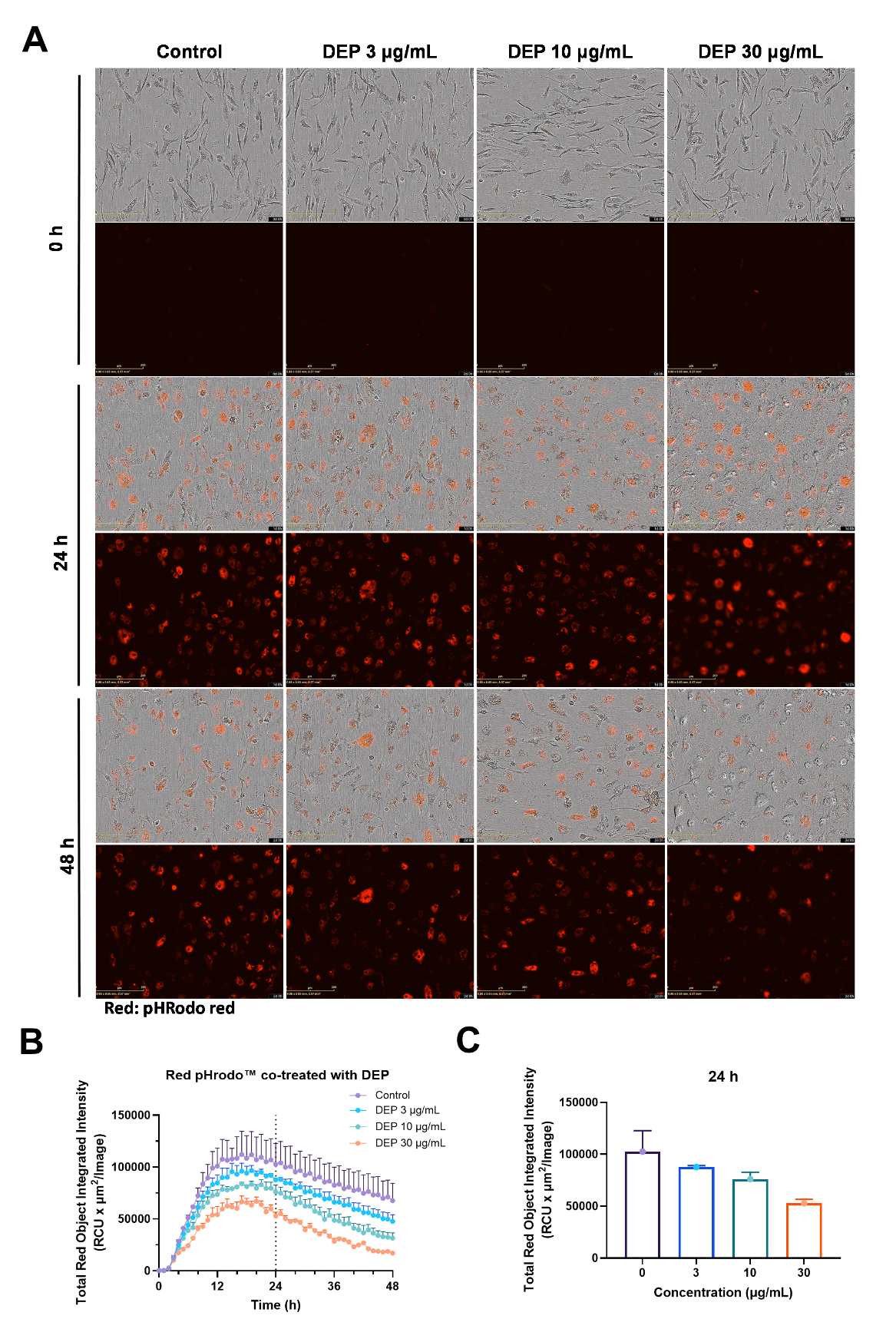
**Supplementary figure S2.** DEP impairs phagocytosis of mouse microglia cells. (**A**) Representative images of primary microglia showing phase contrast and red fluorescence of phagocytosed pHrodo red *E.coli* bioparticles in cells co-treated with solvent or 3, 10, and 30 μg/mL DEP at 0, 12, and 24 h. (**B**) Time-curve of integrated red fluorescence intensity for 3, 10, and 30 μg/mL co-treated microglial cells. (**C**) Bar graph for the integrated fluorescence intensity of pHrodo red *E.coli* bioparticles at 24 h after indicated exposure.

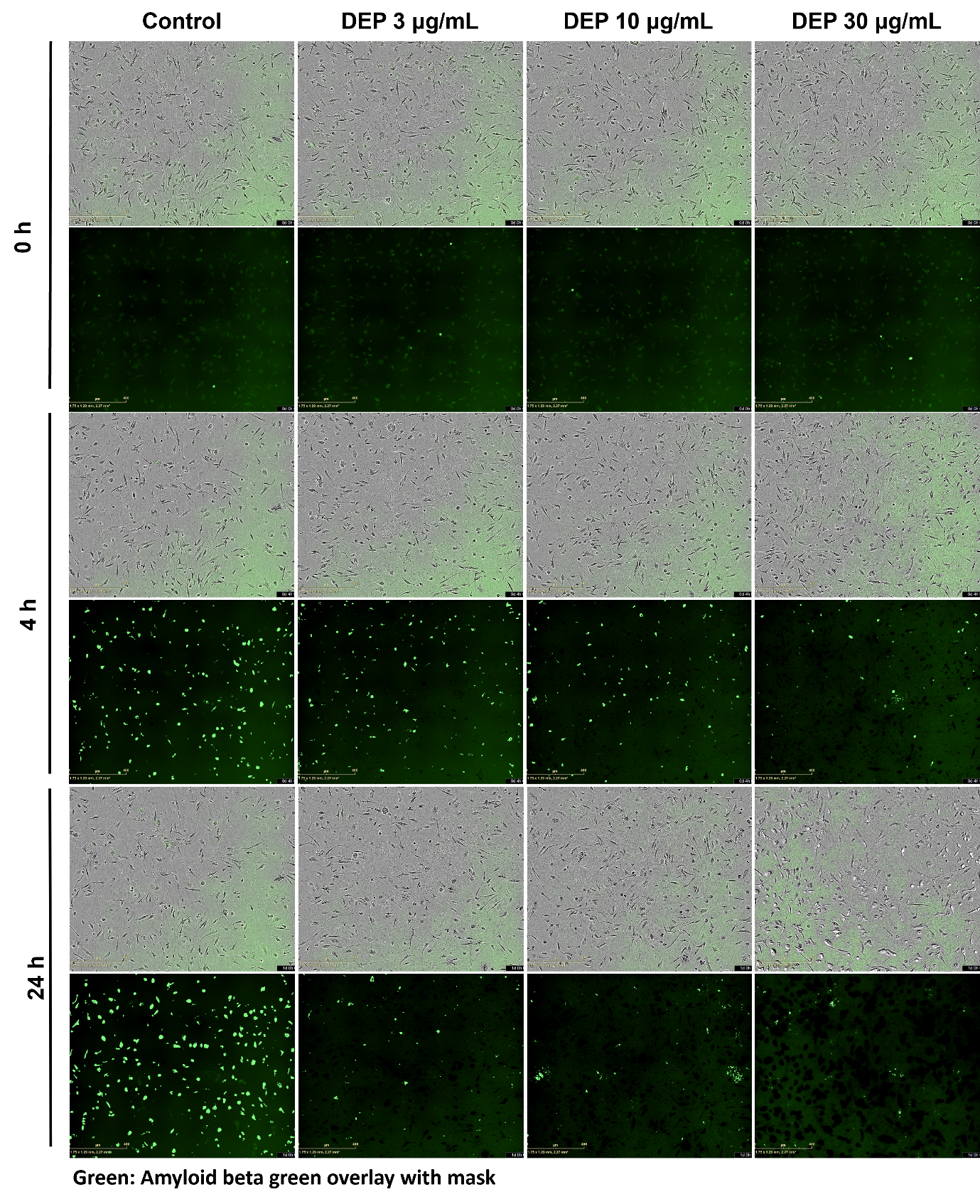
**Supplementary figure S3.** DEP exposure decreases Aβ phagocytosis in mouse microglia cells. (**A**) Representative images of primary microglia showing phase contrast and HiLyte™ Fluor 488-labeled Aβ (1–42, green) after pre-treatment with solvent or 3, 10, and 30 μg/mL DEP at 0, 4, and 24 h. Scale bars 400 μm.

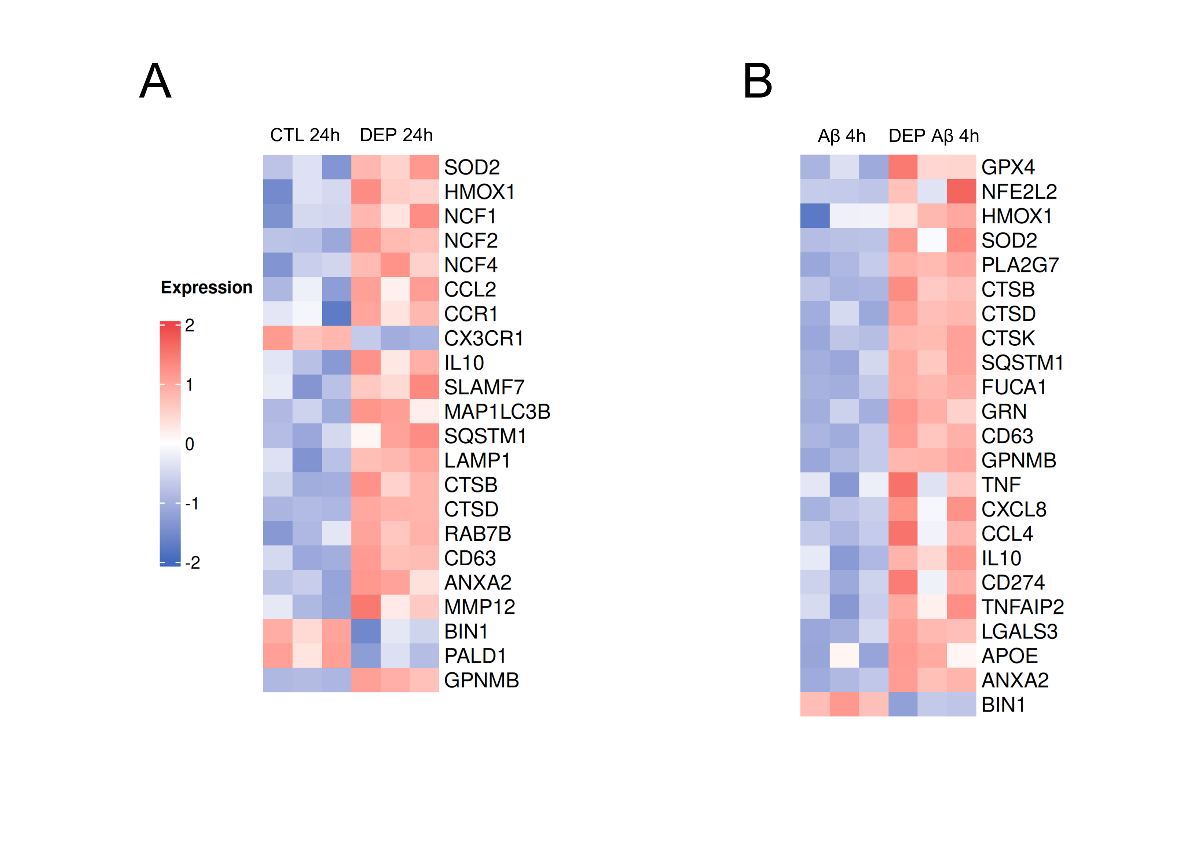

**Supplementary figure S4.** Differential gene expression of iMGL under DEP Exposure and Aβ Challenge. (A) Heatmap showing selected significantly regulated genes in iMGLs following 24 hours of DEP exposure. (B) Heatmap showing selected significantly regulated genes in iMGLs sequentially exposure to DEP for 24 hours followed by a 4-hour Aβ challenge. Significance was determined using an adjusted p-value (padj) < 0.05. n=3 per group.

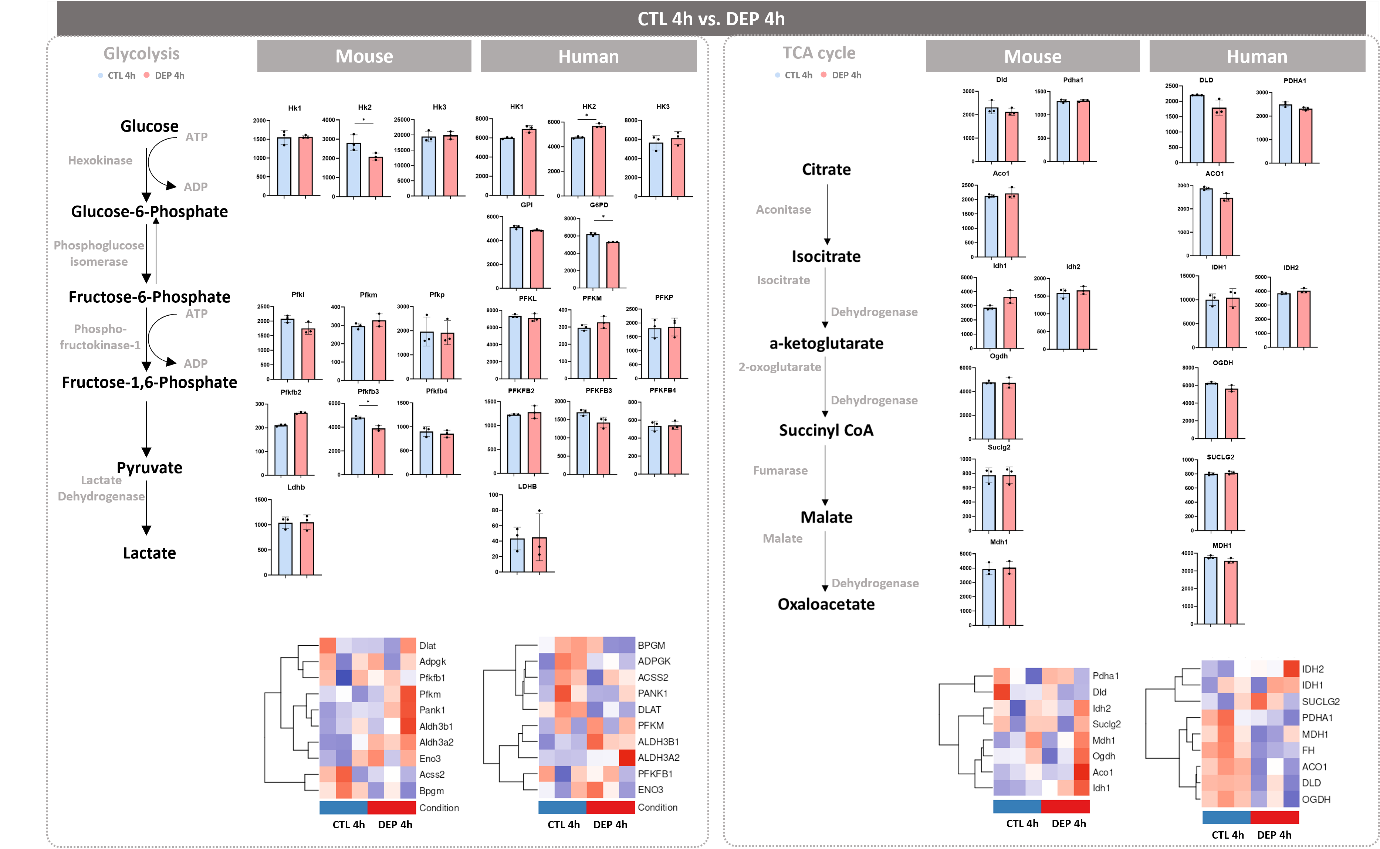

**Supplementary figure S5.** Read count plots and schematic for glycolysis and TCA cycle genes: CTL vs. DEP at 4 h for both mouse microglia and human iMGLs. Control and DEP condition is represented by light blue and light red bars respectively. Data are presented as mean ± SEM. For all comparisons, *p-*value was determined by Wald test with multiple-testing correction by DESEQ2 ^*^*p* < 0.05, ^**^*p* < 0.01, ^***^*p* < 0.001.

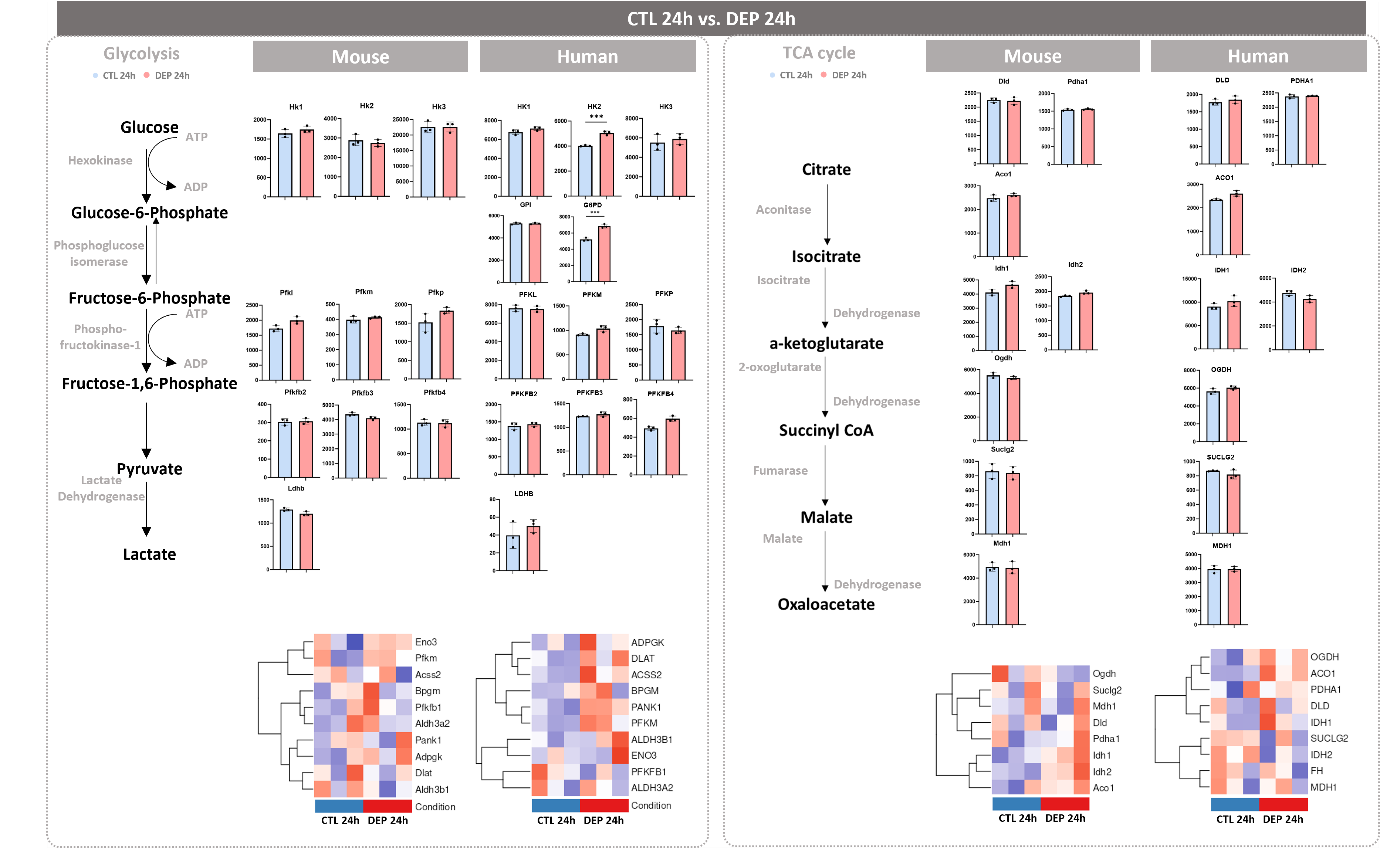

**Supplementary figure S6.**  Read count plots and schematic for glycolysis and TCA cycle genes: CTL vs. DEP at 24 h for both mouse microglia and human iMGLs. Control and DEP condition is represented by light blue and light red bars respectively. Data are presented as mean ± SEM. For all comparisons, *p-*value was determined by Wald–test with multiple-testing correction by DESEQ2. ^*^*p* < 0.05, ^**^*p* < 0.01, ^***^*p* < 0.001.

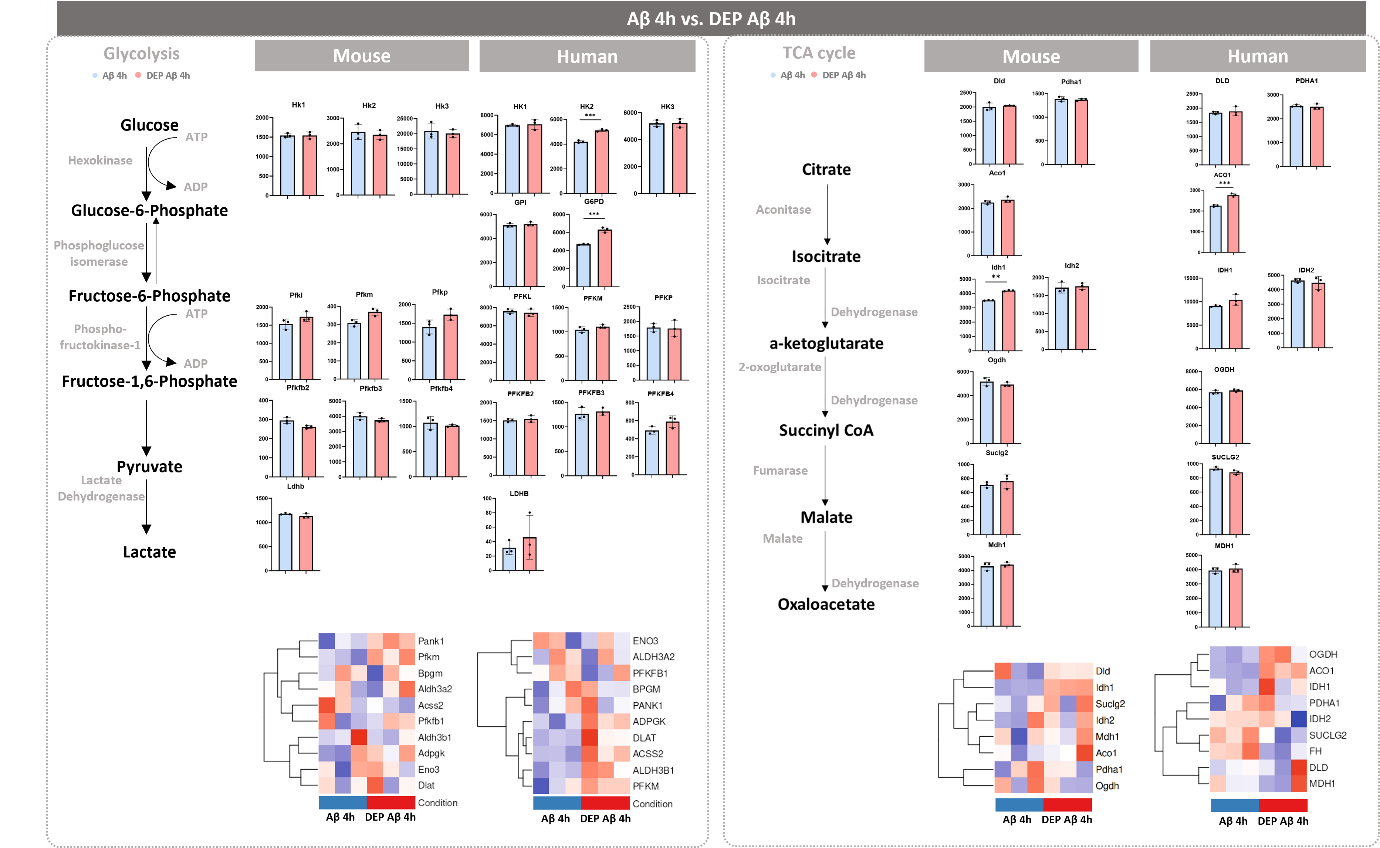

**Supplementary figure S7.** Read count plots and schematic for glycolysis and TCA cycle genes: Aβ 4 h vs. DEP Aβ at 4 h for both mouse microglia and human iMGLs. Control and DEP condition is represented by light blue and light red bars respectively. Data are presented as mean ± SEM. For all comparisons, *p-*value was determined by Wald–test with multiple-testing correction by DESEQ2. ^*^*p* < 0.05, ^**^*p* < 0.01, ^***^*p* < 0.001.
